# Resting state cerebral networks in mouse lemur primates: from multilevel validation to comparison with humans

**DOI:** 10.1101/599423

**Authors:** Clément M. Garin, Nachiket A. Nadkarni, Brigitte Landeau, Gaël Chételat, Jean-Luc Picq, Salma Bougacha, Marc Dhenain

## Abstract

Measures of resting-state functional connectivity allow the description of neuronal networks in humans and provide a window on brain function in normal and pathological conditions. Animal models are critical to further address experimentally the function of brain networks and their roles in pathologies. Here we describe for the first time brain network organization in the mouse lemur (*Microcebus murinus*), a small primate attracting increased attention as a model for neuroscience. Resting-state functional MR images were recorded at 11.7 Tesla. Forty-eight functional regions were identified and used to identify networks using graph theory, dictionary learning and seed-based analyses. Comparison of results issued from these three complementary methods allowed the description of the most robust networks from mouse lemurs. Large scale networks were then identified from resting-state functional MR images of humans using the same method as for lemurs. Strong homologies were outlined between cerebral networks in mouse lemurs and humans.

## 1. Introduction

Blood-oxygen level dependent (BOLD) functional magnetic resonance imaging (fMRI) is largely used to investigate brain function in response to specific tasks. In the absence of explicit tasks (*i.e.* in resting state conditions) patterns of oscillations of the fMRI signal are similar in functionally connected brain structures (Biswal et al., 1995). The detection of the synchronicity of BOLD signal in various brain regions in resting state conditions can thus be used to describe cerebral network organization. In particular this allows the characterization of *i.* local regions in which highly coordinated neuronal activity occurs and *ii.* large scale networks composed of widespread functional regions connected together (Biswal et al., 1995; Power et al., 2014).

Studies of brain networks have contributed to many breakthroughs in the understanding of brain function in normal as well as in pathological conditions such as Alzheimer’s or Parkinson’s diseases (Buckner et al., 2005; Gao and Wu, 2016). However, many questions remain concerning both the technique and interpretation of resting state fMRI. For example, both the role of resting state networks in cerebral function, and the biological mechanisms underlying their activity, are still partly unknown. Also, how their modulations impact behaviour and cognition in pathological conditions is still debated (Mohan et al., 2016).

Using animal models is critical to further address these questions. Indeed, in animals it is possible to artificially stimulate neuronal activity to characterize biological mechanisms underlying network function (Gerits et al., 2012). Another interest of studying neuronal networks in animals is to evaluate how evolution has driven network architecture and to assess to what extent animal behaviours and ecology (Burkart et al., 2016) have impacted this architecture. Finally, animals can be used to model diseases and explore the impact of pathological processes on brain networks.

Various analysis pipelines have been proposed to investigate neuronal networks in humans and animals. For example, large scale networks were identified using data-driven methods relying on spatial map decomposition (dictionary learning (Varoquaux et al., 2011), independent component analysis (Damoiseaux et al., 2006)) or on graph theory (modularity analysis (Grayson et al., 2016)), as well as hypothesis-driven methods (seed-based analysis (Hutchison et al., 2014)). These methods are based on different algorithms and each one has its own inherent advantages and disadvantages (Lee et al., 2013). They can provide complementary approaches for identifying networks in unexplored animals.

The mouse lemur (*Microcebus murinus*) is a primate attracting increased attention in neuroscience research. This small animal (typical length 12cm, 60-120g weight) is arboreal and nocturnal. It has a decade-long lifespan and is a model for studying cerebral aging (Sawiak et al., 2014) and various diseases such as diabetes-related encephalopathy (Djelti et al., 2016), Parkinson’s disease (Mestre-Frances et al., 2018), or Alzheimer’s disease (Kraska et al., 2011). It has a key position on phylogenetic trees of primates and is used to investigate primate brain evolution (Montgomery et al., 2010). Characterizing its cerebral networks is thus useful in the context of comparative biology as well as for further use of this animal to model various pathologies. Thus, the first aim of this study was to characterize neuronal networks in mouse lemurs. Our second objective was to implement a protocol that could define functional regions directly from resting-state fMR images and to compare large scale networks identified with data-driven and hypothesis-driven methods to assess the robustness of the identified networks. Our third objective was to compare resting state networks identified in lemurs with those identified in humans using the same procedure.

Resting state functional MR images were recorded from 14 mouse lemurs at 11.7 Tesla. These images enabled the identification of 48 functional regions using dictionary learning (Varoquaux et al., 2011). These regions were concatenated into a 3D functional atlas covering most of the brain and were used as nodes for whole brain network characterization. Large scale networks were identified using several methods based on graph theory, dictionary learning and seed-based analysis. They included default-mode-like, visual, fronto-temporal, somato-motor, basal ganglia and thalamic networks. These networks were then compared to large scale networks in humans. We found a strong homology between cerebral networks in mouse lemurs and humans.

## 2. Results

### 2.1. Identification of local functional regions and concatenation in a 3D functional atlas

Resting state fMR images were recorded from 14 anaesthetised (isoflurane 1.25-1.5%) mouse lemurs at 11.7 Tesla (Suppl. Table 1). Images were recorded using a gradient-echo echo planar imaging (EPI) sequence. Each animal was scanned twice with an interval of 6 months.

Organisation of whole brain networks can be modelled using graph theory. During this modelling, whole brain networks are defined as a set of nodes (basic elements of the system) and edges (allowing relationships between nodes). The identification of nodes can be based on the use of anatomical atlases (Ghahremani et al., 2016) or on the use of study-specific functional atlases that identify local functional regions (Ma et al., 2018).

Here, we identified local functional regions by performing a dictionary learning based on a large number of sparse components (SCs). This method extracts maps of cerebral networks from fMRI data and relies on sparsity-based decomposition of the signal. Multi-animal dictionary learning analyses of resting state fMR images were performed in mouse lemurs using 35 components (Fig. 1). Each component was manually classified using anatomical (Bons et al., 1998; Nadkarni et al., 2018) and Brodmann atlases (Brodmann, 1999 (original in 1909); Le Gros Clark, 1931). First, brain regions were classified based on their locations within the frontal, parietal, temporal and occipital lobes as well as subcortical and midbrain regions. The 35 components were used to create a 3D functional atlas of the brain (Fig. 2). Some single components were associated to bilateral structures as shown, for example, for the precentral cortex in Fig. 1. These bilateral regions were classified as two different regions (*i.e.* one in each hemisphere). Thus, 48 local functional regions (27 cortical, 21 subcortical) could be extracted from the 35 component dictionary analysis (Table 1). They can be downloaded from https://www.nitrc.org/projects/fmri_mouselemur/.

**Figure 1.**
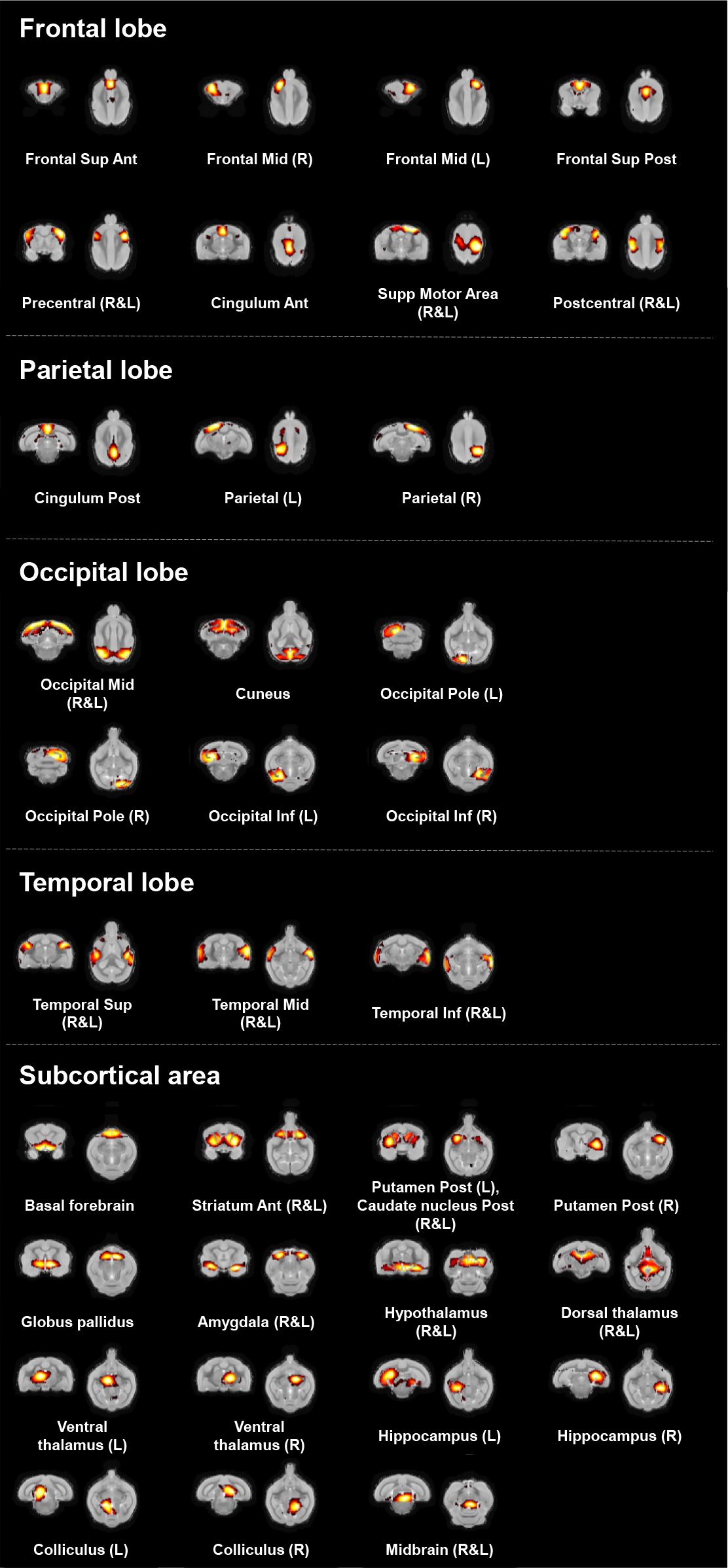
Regions of functional activity identified in mouse lemurs. Regions of functional activity were identified following dictionary learning analyses of resting state fMR images using 35 components. They are shown on coronal and axial anatomical templates with an automatic slice selection based on the center of mass of each component. All components were organized within five anatomical areas: frontal, parietal, occipital, temporal, and subcortical regions.

**Figure 2.**
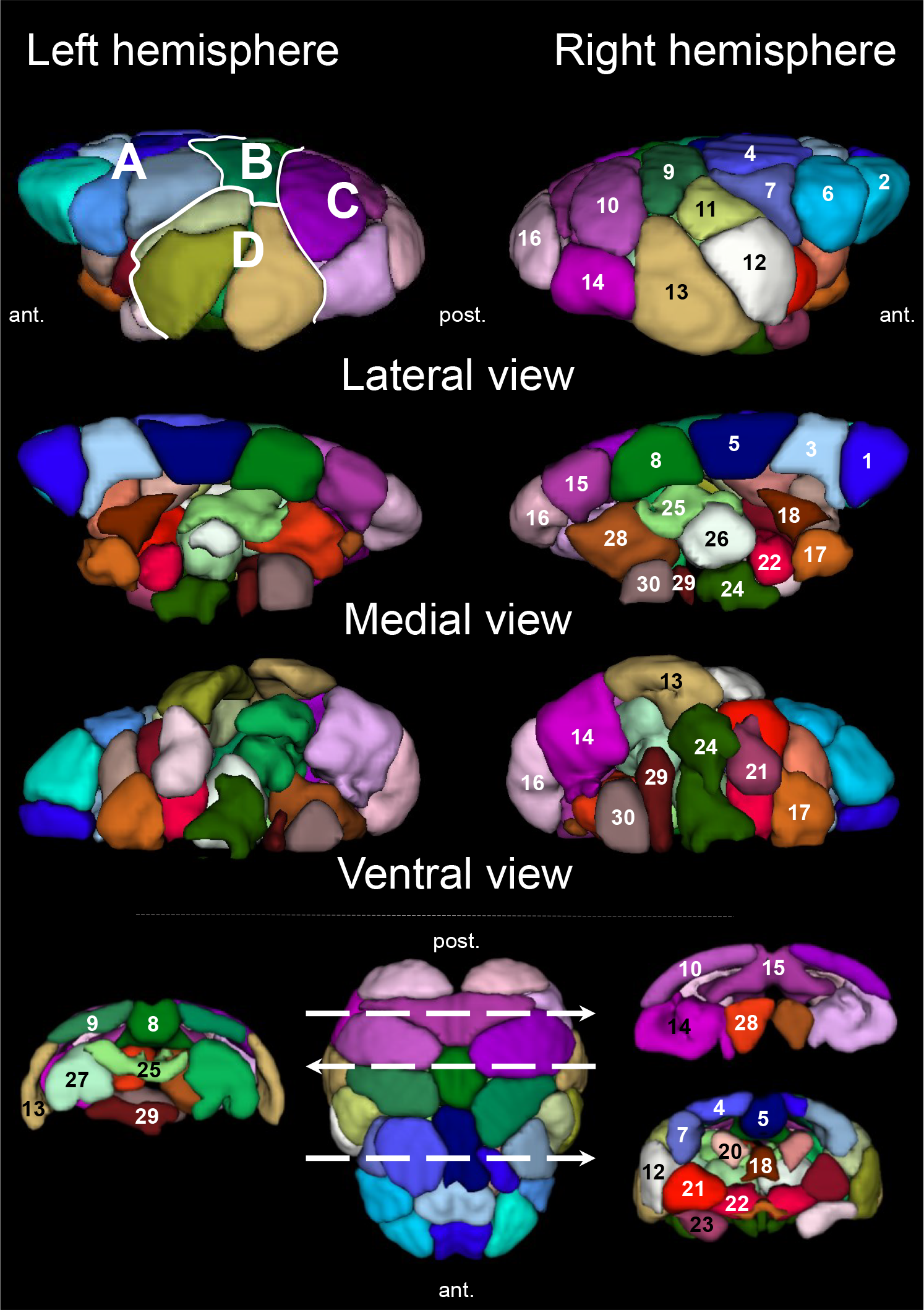
Mouse lemur 3D functional atlas based on dictionary learning. Forty eight local functional regions were identified following dictionary learning analyses of resting state fMR images using 35 components. Brain regions were classified based on their locations within the frontal (A), parietal (B), occipital (C), and temporal (D) lobes. We display three different views and three slices extracted from the functional atlas. 1. Frontal Superior Anterior, 2. Frontal Middle, 3. Frontal Superior Posterior, 4. Supplementary Motor Area, 5. Cingulum Anterior, 6. Precentral, 7. Postcentral, 8. Cingulum Posterior, 9. Parietal, 10. Occipital Middle, 11. Temporal Superior, 12. Temporal Middle, 13. Temporal Inferior, 14. Occipital Inferior, 15. Cuneus, 16. Occipital Pole, 17. Basal forebrain, 18. Septal nuclei, 19. Striatum Anterior, 20. Caudate nucleus Posterior, 21. Putamen Posterior, 22. Globus pallidus, 23. Amygdala, 24. Hypothalamus, 25. Dorsal thalamus, 26. Ventral thalamus, 27. Hippocampus, 28. Colliculus, 29. Pons, 30. Midbrain.

**Table 1.**
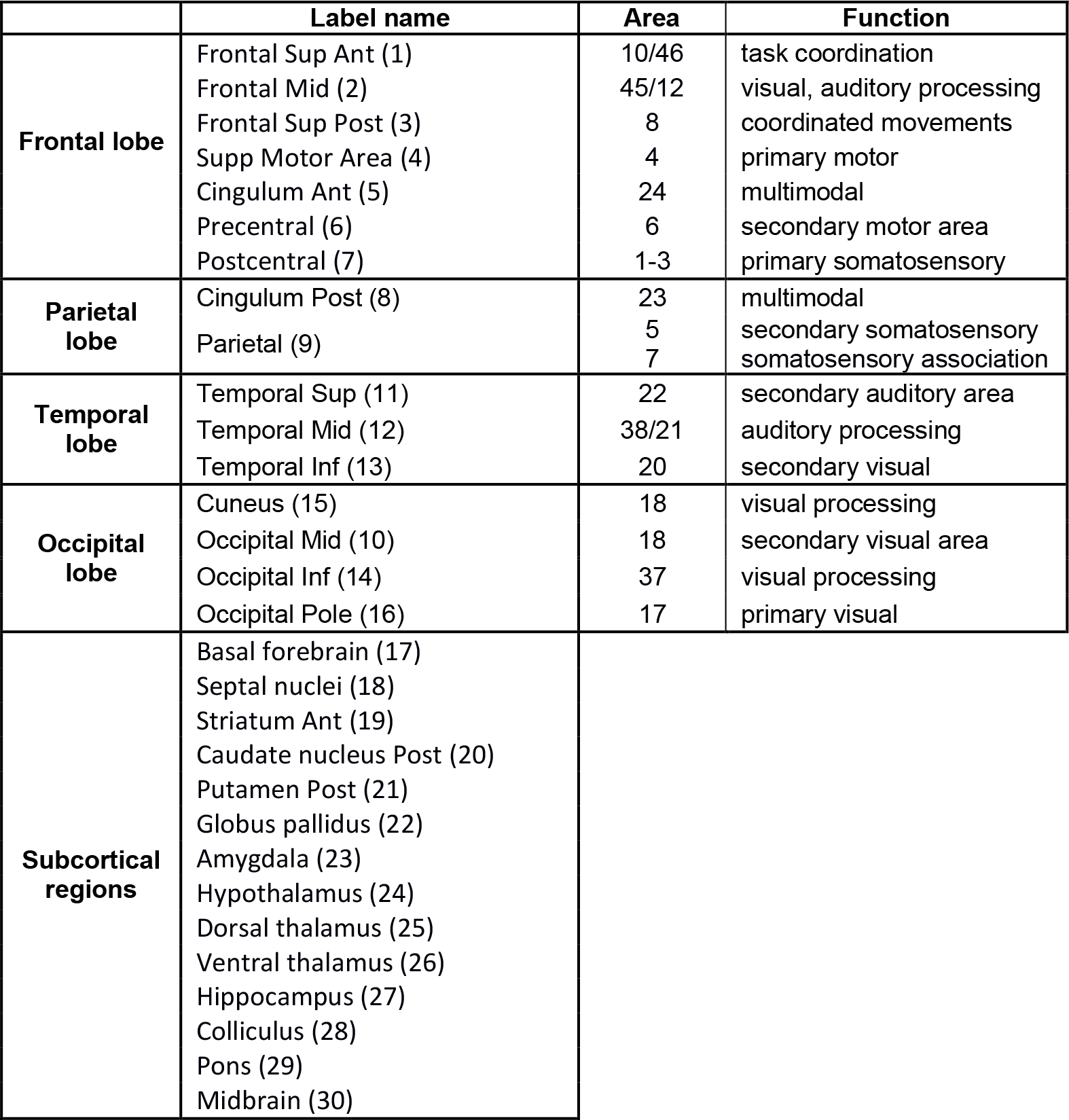
Identification of functional regions of the mouse lemur brain. Brain regions were classified based on their locations within the frontal, parietal, temporal, or occipital lobes as well as subcortical regions. Each labelled region was compared to cytoarchitectonic (Brodmann, 1999 (original in 1909); Le Gros Clark, 1931) and anatomical atlases of the mouse lemur (Bons et al., 1998; Nadkarni et al., 2018) and of the human “AAL for SPM12” atlas (Tzourio-Mazoyer et al., 2002) to evaluate the Brodmann areas that were the closest to the identified regions. A function is also proposed for each region following expectations from Brodmann classification.

|  | Label name | Area | Function |
| --- | --- | --- | --- |
| **Frontal lobe** | Frontal Sup Ant (1) | 10/46 | task coordination |
|  | Frontal Mid (2) | 45/12 | visual, auditory processing |
|  | Frontal Sup Post (3) | 8 | coordinated movements |
|  | Supp Motor Area (4) | 4 | primary motor |
|  | Cingulum Ant (5) | 24 | multimodal |
|  | Precentral (6) | 6 | secondary motor area |
|  | Postcentral (7) | 1-3 | primary somatosensory |
| **Parietal lobe** | Cingulum Post (8) | 23 | multimodal |
|  | Parietal (9) | 5<br>7 | secondary somatosensory<br>somatosensory association |
| **Temporal lobe** | Temporal Sup (11) | 22 | secondary auditory area |
|  | Temporal Mid (12) | 38/21 | auditory processing |
|  | Temporal Inf (13) | 20 | secondary visual |
| **Occipital lobe** | Cuneus (15) | 18 | visual processing |
|  | Occipital Mid (10) | 18 | secondary visual area |
|  | Occipital Inf (14) | 37 | visual processing |
|  | Occipital Pole (16) | 17 | primary visual |
| **Subcortical regions** | Basal forebrain (17)<br>Septal nuclei (18)<br>Striatum Ant (19)<br>Caudate nucleus Post (20)<br>Putamen Post (21)<br>Globus pallidus (22)<br>Amygdala (23)<br>Hypothalamus (24)<br>Dorsal thalamus (25)<br>Ventral thalamus (26)<br>Hippocampus (27)<br>Colliculus (28)<br>Pons (29)<br>Midbrain (30) |  |  |

### 2.2. Large scale brain networks in mouse lemurs

The quantification of correlations of temporal evolution of BOLD fMRI signal between two regions (or nodes) provides an index of the “functional connectivity” between these nodes. Here, the 48 functional regions identified with the dictionary learning analysis were used as nodes for graph analysis of the mouse lemur brain. A 3D-view of the mouse lemur network based on these 48 functional regions is presented in Suppl. Fig. 1. Partial correlation matrices were created using fully preprocessed MR images by calculating the partial correlation coefficients between temporal evolutions of BOLD MR signals within each region of this 3D functional network.

#### 2.2.1. Modularity and large scale network identification based on graph analysis

In graph theory, large scale networks are defined as community structure (or modules), which are groups of nodes connected densely and sparsely with nodes from other modules. The modularity of a partition (Q) is the degree to which a network can be subdivided into non-overlapping groups of nodes with maximum within-group connections and minimum number of between-group connections (D. B. Vincent et al., 2008). Here, the average partial correlation matrix was used to evaluate the modular structure of the mouse lemur brain by graph theory. Q was calculated to assess the ability of this weighted undirected matrix to be segregated into non-overlapping groups of nodes. A high modularity value (Q = 0.43) was obtained which suggests a prominent modular structure of mouse lemur brain networks. This modularity index was associated with the classification of the matrix into 6 modules (large scale networks) (Fig. 3, Suppl Table 2). Each functional region was associated with one and only one network. These networks were identified as:

**Figure 3.**
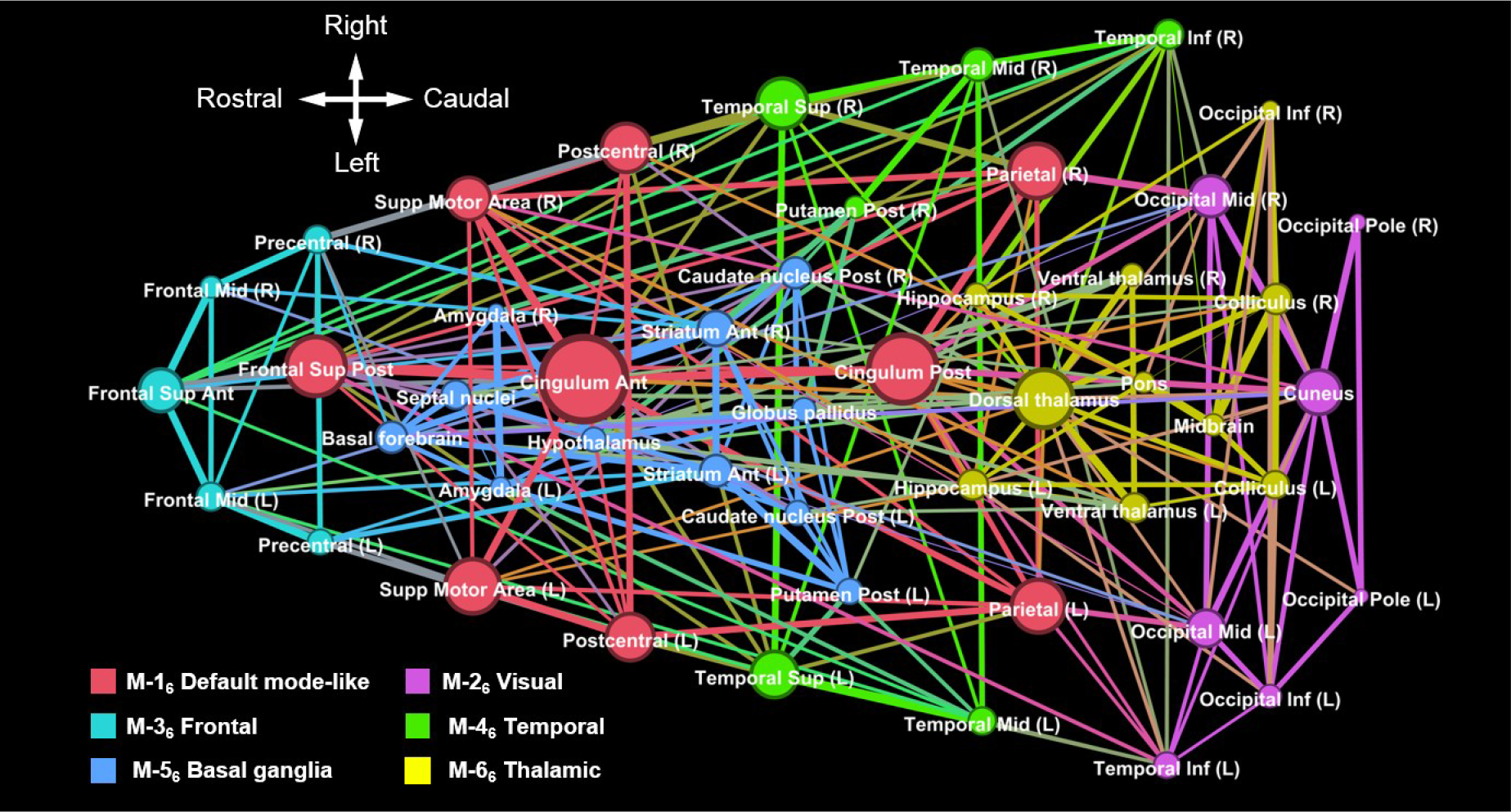
Mouse lemur networks identified using graph analysis based on 48 functional regions. Using graph analysis, we partitioned the mouse lemur brain into six cortical and subcortical modules. A color and a name were assigned to each module. Colors highlight interactions between different nodes, *i.e.* they outline large scale networks. Eigenvector centrality, a measure of node influence, is represented by the node size.

M-1_6_ – Default mode network-like (DMN-like). This module involved posterior and anterior cingulum, superior posterior frontal and parietal cortices. In other species, these regions are reported to be part of the DMN (Belcher et al., 2013; Hutchison et al., 2010; J. L. Vincent et al., 2007). This module also embedded nodes from the superior motor area and postcentral cortices.

M-2_6_ – Visual. This module involved the cuneus, the occipital pole, the middle, the inferior occipital and the inferior temporal cortices. Those clusters correspond to visual areas and regions involved in integration of visual information.

M-3_6_ – Frontal. This module involved nodes from frontal and precentral cortices.

M-4_6_ – Temporal. This module embedded temporal structures usually implicated in response to auditory stimuli as well as the right posterior putamen.

M-5_6_ – Basal ganglia. This module embedded the anterior striatum, the posterior striatum (posterior caudate nucleus and posterior putamen), the amygdala, basal forebrain, septal nuclei, as well as the hypothalamus and globus pallidus.

M-6_6_ – Thalamic. This network involved a large number of subcortical regions including and surrounding the thalamus, the hippocampus, the colliculi and the midbrain.

#### 2.2.2. Identification of large scale networks based on dictionary learning

We then wondered whether the six previously identified modules could be identified with dictionary learning analysis, another data-driven method. A six-component analysis revealed bilateral networks spread over the whole brain (Fig. 4, Suppl. Table 2). Four networks (the DMN, visual, basal ganglia and thalamic) were very similar to those identified with the module analysis. One network (fronto-temporal) was a concatenation of two networks identified by module analysis. The last network (somato-motor) was not identified with module analysis. Unlike for the graph analysis some functional regions (*e.g.* the anterior cingulate cortex) could be attributed to different networks (*e.g.* the DMN, fronto-temporal and somato-motor networks). More precisely, the networks were identified as:

**Figure 4.**
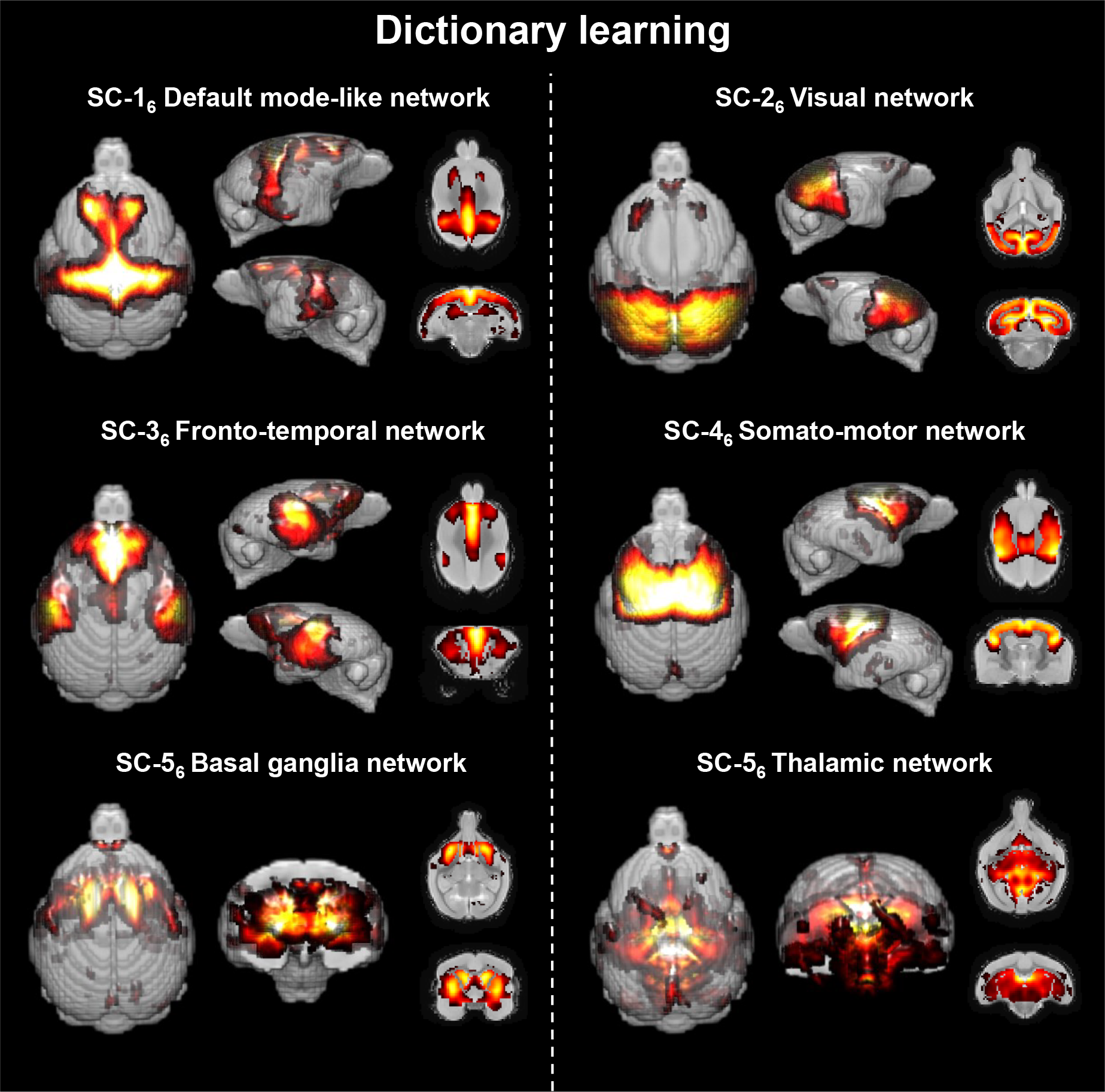
Cerebral networks identified following six component dictionary learning in mouse lemurs. This analysis revealed bilateral networks that included several regions spread over the whole brain classified as default mode-like, visual, fronto-temporal, somato-motor, basal ganglia and thalamic networks.

SC-1_6_ – DMN. This network involved structures identified with graph analysis (posterior and anterior cingulum cortices, superior posterior frontal and parietal cortices). Some nodes (superior motor area and postcentral cortices) identified as part of the DMN by graph analysis were not detected with dictionary learning.

SC-2_6_ – Visual. This network involved the same nodes as those detected with module analysis (occipital pole, middle, inferior occipital and inferior temporal cortices), except the inferior temporal cortex.

SC-3_6_ – Fronto-temporal. This network involved several regions that were identified as frontal or temporal network with graph analysis. It also included the anterior cingulum cortex.

SC-4_6_ – Somato-motor. This network embedded frontal and parietal regions located above the Sylvian fissure (corresponding to Brodmann 1-3 (primary region involved in body sensation), 4 (primary motor region) and 6 (secondary motor region)) and temporal regions surrounding the Sylvian fissure. This network could thus be involved in somato-motor activities.

SC-5_6_ – Basal ganglia. This network involved the same regions as those identified for this network with module analysis except for the hypothalamus and globus pallidus.

SC-6_6_ – Thalamic. This last network involved mostly the same regions as the ones identified with graph analysis. In addition, it included the basal forebrain, septal nuclei and globus pallidus.

#### 2.2.3. Identification of large scale networks based on seed-based analysis

Another way to analyse cerebral networks is to perform seed-based studies. This method evaluates the relationships between mean BOLD signal in a brain region (seed) and BOLD signal in any voxel of the brain. Here, the seeds corresponded to the 48 previously identified functional regions. Some seeds were only connected with voxels from the same brain region and were not further explored (*i.e.* the visual and thalamic networks, SB-2_6_ and SB-6_6_ in Fig. 5). Four seeds were connected with voxels localized in brain networks previously described with the graph analysis and dictionary learning methods (*i.e.* the DMN, fronto-temporal, somato-motor and basal ganglia networks, Fig. 5). Two networks identified with other methods were not identified by seed-based analysis (the visual and thalamic networks). As for dictionary learning, some structures (*i.e.* the anterior cingulum cortex) could be attributed to different networks (Suppl. Table 2). More precisely, the networks highlighted by seed-based analysis are described as follows.

**Figure 5.**
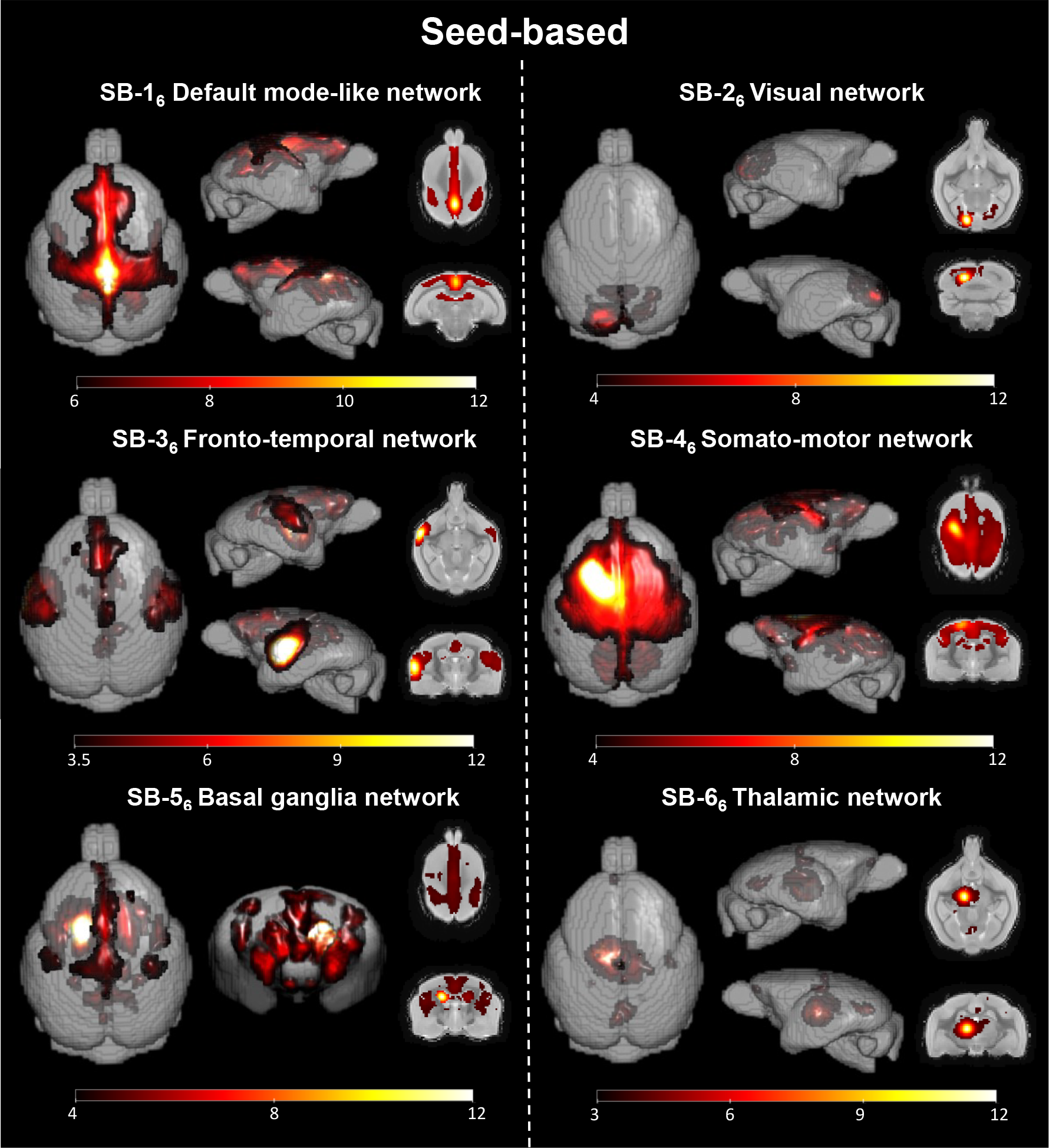
Mouse lemur resting-state networks characterized with seed-based analysis. Each image highlights mean *z*-statistic maps of regions connected to a cerebral seed. Seed-based analysis detected four of the six previous large scale networks identified with dictionary learning: default mode-like, fronto-temporal, somato-motor, and basal ganglia (seeds positioned in the posterior cingulate cortex, the left medial temporal cortex, the left superior frontal cortex and the left posterior caudate nucleus, respectively). Visual and thalamic networks that were detected with dictionary learning were not detected with seed-based analysis: SB-2_6_ and SB-6_6_ display lack of large network detection using seeds in the left occipital cortex and the left ventral thalamus. Color bars represent *z*-statistic values.

SB-1_6_ – DMN. The seed from the posterior cingulum cortex (PCC) is usually used to define the DMN. Here, using this seed we highlighted highly connected voxels in the regions identified as DMN with graph analysis and dictionary learning methods (posterior and anterior cingulum cortices, superior posterior frontal and parietal cortices). Additional parts of this network were also identified (middle frontal cortex and dorsal thalamus).

SB-3_6_ – Fronto-temporal. The seed from the left middle temporal cortex was connected with the right middle and superior temporal cortices, superior anterior frontal cortex, superior posterior frontal cortex and anterior cingulum cortex.

SB-4_6_ – Somato-motor. Using a seed in the left superior motor area, we highlighted a network englobing several regions included in the somato-motor network identified by dictionary learning (fronto and parietal cortices, superior temporal regions, anterior cingulum cortex). Voxels from the middle frontal, superior posterior frontal cortex, posterior cingulum cortices as well as the posterior caudate nucleus and dorsal thalamus were also associated with this network.

SB-5_6_ – Basal ganglia. Using the posterior caudate nucleus (left) as a seed, we highlighted a basal ganglia network that involved the striatum. It was already identified for this network with graph analysis and dictionary learning. Voxels from the superior posterior frontal cortex and anterior cingulum cortices were also associated with this network.

### 2.3. Functional hubs and small-worldness features of mouse lemur brains

#### 2.3.1. Brain hubs in mouse lemurs

Whole brain networks can also be characterized using various descriptors. One of these descriptors, "hubness", describes the centrality of nodes in the network. This is a measure of node influence within the whole brain network. It can be measured by eigenvector centrality. For each node, this index is mainly calculated based on its partial correlation values (edges) with all regions of the 3D functional atlas, weighted by the eigenvector scores of its neighbourhood nodes. In other words, nodes which display high eigenvector centrality scores are strongly linked to other nodes and/or to strongly connected nodes. Here, eigenvectors were presented as histograms (Fig. 6) or as the size of the nodes in the graphical representation of the networks (Fig. 3). The 3 nodes presenting the highest eigenvector centrality were the anterior cingulum cortex, the posterior cingulum cortex, and the superior posterior frontal cortex. These three regions belong to the DMN. The dorsal thalamus was the next region showing highest hubness properties. Then the following hubs involved the parietal cortex, superior motor area, as well as the superior temporal and postcentral cortices (Fig. 6).

**Figure 6.**
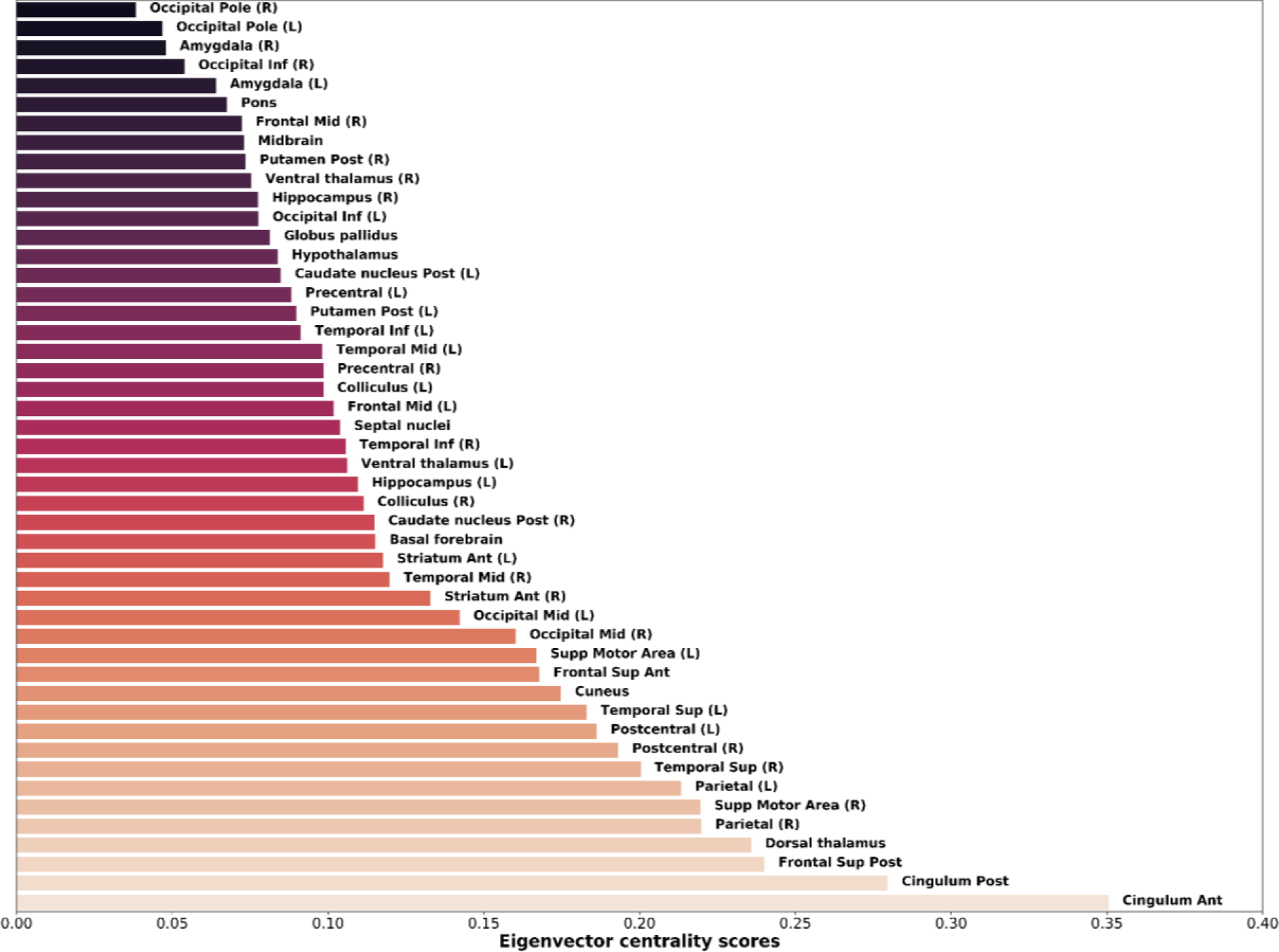
Eigenvalue centrality scores, reflecting "hubness", in mouse lemur brain regions. The three regions displaying the highest scores were the anterior cingulate cortex, the posterior cingulate cortex and the central frontal cortex. The dorsal thalamus was the next region showing highest hubness properties. Then the following hubs involved the parietal cortex, superior motor area, as well as the superior temporal and postcentral cortices.

#### 2.3.2. Small-worldness of mouse lemur brain networks

Network topology describes properties of regional specialization and global information transfer efficacy. It can be classified into three main classes: random, lattice and small-world networks (Telesford et al., 2011). Network topology can be characterized using two small-world coefficients (σ and ω) (NetworkX (Hagberg et al., 2008)). Small-world networks have σ values superior to 1 and ω values close to 0 (Telesford et al., 2011). In mouse lemurs these coefficients (σ = 1.47 and ω = 0.39) indicated small-world properties. Usually, mammal brains have small-worldness topology (Mechling et al., 2014).

### 2.4. Cerebral networks in humans

We then wondered how comparable mouse lemur and human brain networks are. To answer to this question, resting state fMRI data were recorded from 42 healthy humans ranging from 41 to 60 years old at 3.0 Tesla using an interleaved 2D T2* SENSE EPI. Participants were asked to keep their eyes closed and relax without falling asleep during image acquisition. Human images were then processed with the same graph analysis and dictionary learning algorithms as mouse lemur images.

Local functional regions were identified using a dictionary learning based on 35 components. Single components spread on bilateral structures were dissociated into two different regions (*i.e.* one in each hemisphere). Ultimately, the brain was partitioned into 56 local functional regions (55 cortical, 1 subcortical). They were named based on the “AAL for SPM12” atlas (Tzourio-Mazoyer et al., 2002) (Suppl. Fig. 2)

As for mouse lemurs, the 56 functional regions identified with the dictionary learning analysis were used as nodes for large scale network analysis. First, we calculated partial correlation coefficients between temporal evolutions of BOLD MR signals within each region of the 3D functional atlas. The obtained correlation matrix was used to calculate the matrix modularity value (Q = 0.56). This index was associated with the segregation of the matrix into 6 modules that were classified as default mode, visual, frontal, temporal somato-motor, and temporo-insular networks (Suppl. Fig. 3).

Then large scale networks were further characterized in humans using a dictionary learning analysis with 6 components (Fig. 7, Table 2). The 6 networks identified could be classified as the default mode, visual, fronto-supramarginal (classified as control-executive network in (Solé-Padullés et al., 2016)), somato-motor, temporal, and a fronto-parietal network (classified as attention network in (Raichle, 2011)).

**Figure 7.**
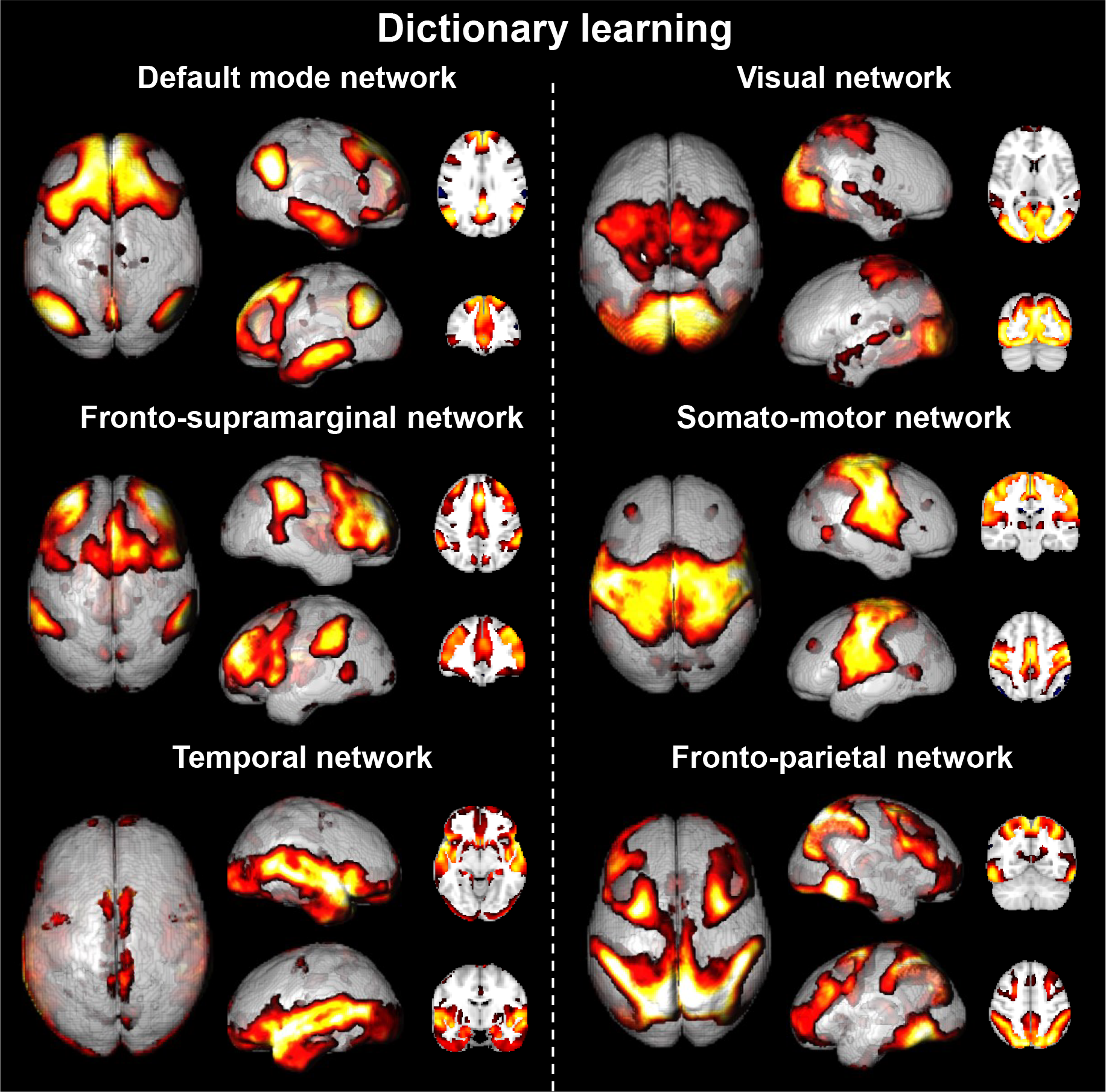
Human cerebral networks identified following six component dictionary learning. The spatial map decomposition extracted 6 cortical networks commonly observed in the literature (DMN, visual, fronto-supramarginal, somato-motor, temporal, fronto-parietal). This analysis was performed with similar pretreatments as for the mouse lemurs.

**Table 2.**
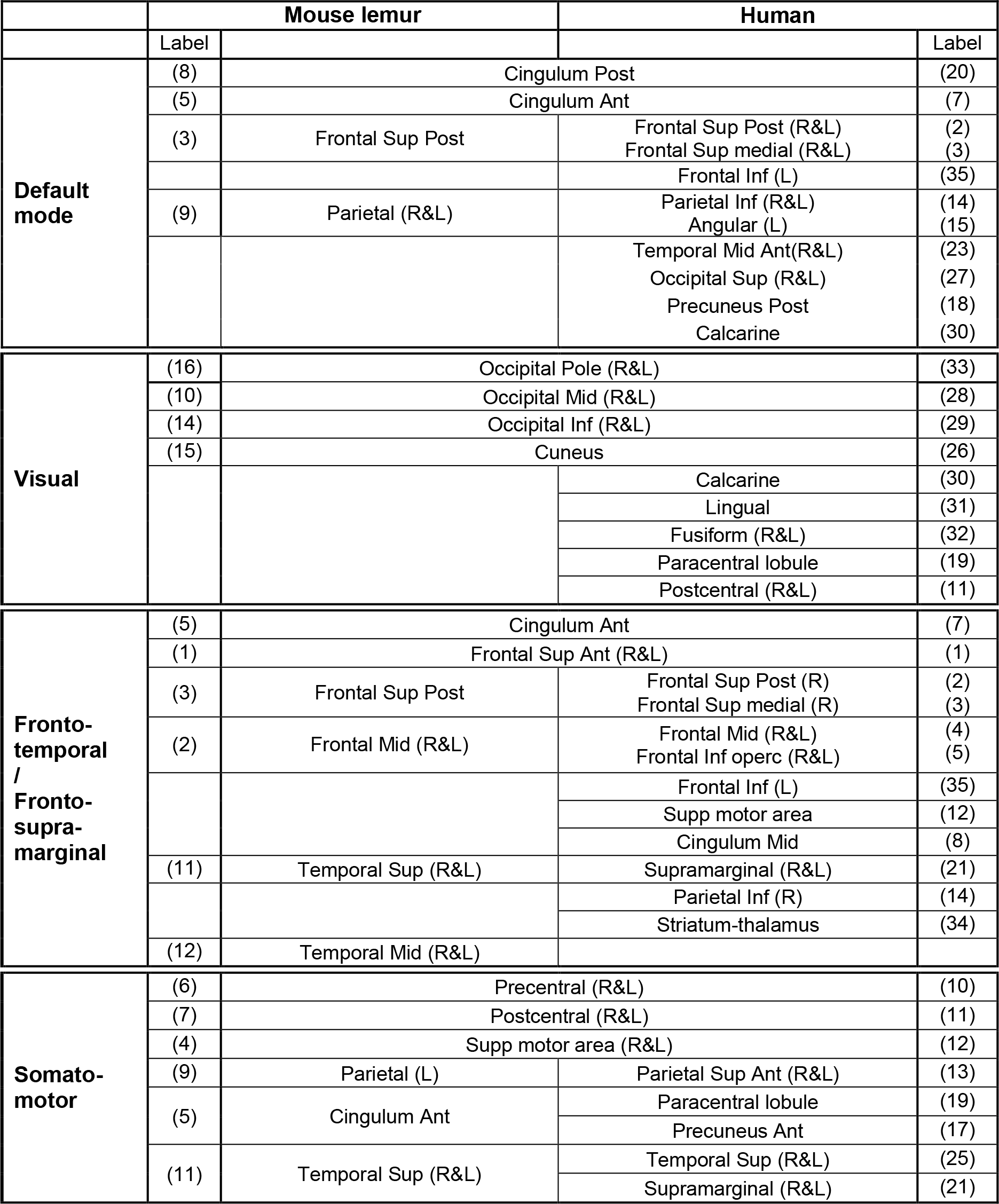
Comparison of the regions belonging to the different networks extracted in mouse lemurs and humans. Regions that were identified with different methods are grouped within a single case. The 3D functional atlas of each species was pasted on different networks obtained by dictionary learning. A region was considered to belong to a network when more than 30% of its volume belonged to this network. The fit between two regions with different names was based on the anatomical proximity. Labels represent the number corresponding to this region in Figure 2 for lemurs and Supplementary Figure 2 for humans.

### 2.5. Functional hubs and small-worldness features of human brains

Eigenvector centrality and network topology were evaluated in humans using the same procedures as for mouse lemurs. Eigenvector centrality was presented as histograms (Fig. 8) or as the size of the nodes in the graphical representation of the networks (Suppl. Fig. 3). The 3 nodes presenting the highest eigenvector centrality were the parietal inferior (right and left) and the precuneus posterior. Then the next hubs were located in the middle frontal cortex (left), the angular region (left) and the posterior cingulum cortex. All these regions except the middle frontal cortex belong to the DMN. Regarding network topology, as expected we found small-world properties in the human brain (σ = 1.1 and ω = 0.08).

**Figure 8.**
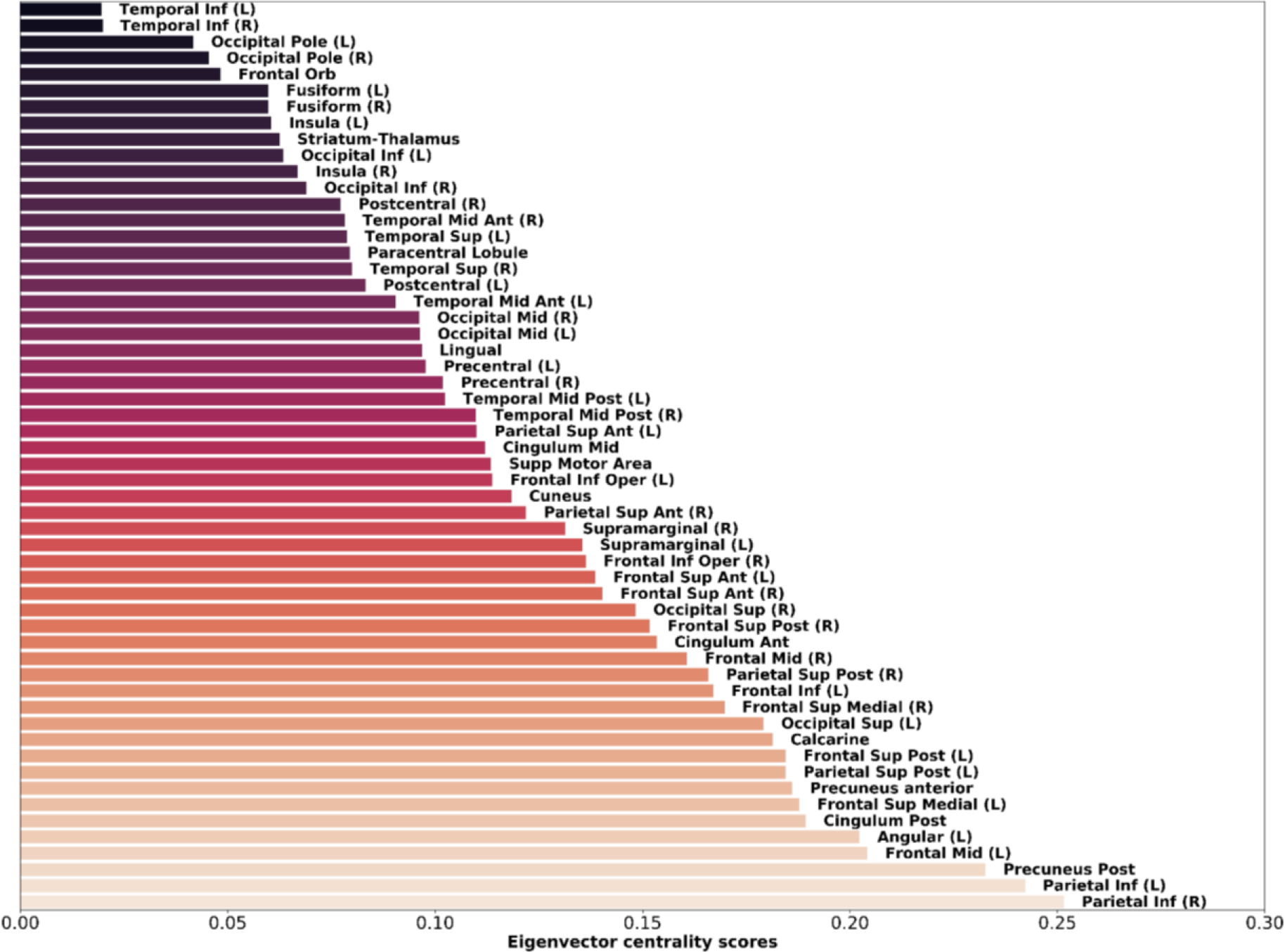
Eigenvalue centrality scores, reflecting "hubness", in human brain regions. The 3 nodes presenting the highest eigenvector centrality were the parietal inferior (right and left) and the precuneus posterior. Then the next hubs were located in the middle frontal cortex (left), the angular region (left) and the posterior cingulum cortex.

## 3. Discussion

This study provides a detailed characterisation of the organisation of functional networks in mouse lemur primates under isoflurane sedation. Complementary analyses based on dictionary learning, seed-based studies and graph analysis highlighted 48 local functional regions that could be grouped into several large scale networks. We also identified the main hubs and small-world characteristics of mouse lemur brains. Human brain networks were also analysed with algorithms similar to those used in lemurs in order to compare networks in both species.

### 3.1. Parcellation of functional regions within mouse lemur brains

Up to now, description of mouse lemur functional organisation was based on cytoarchitectonic atlases (Bons et al., 1998; Le Gros Clark, 1931; Nadkarni et al., 2018). Here, using dictionary learning with a large number of components, we created a 3D map of 48 local functional regions. The quality of this functional atlas was supported by the bilateralism of the extracted regions. One of the strengths of this functional map is that it can be used to create a whole brain graph that relies on brain function rather than on anatomical boundaries. Studies of animal resting state networks often used regions of interest based on anatomical atlases (Li and Zhang, 2018), as opposed to functional atlases. The latter approach is preferable since anatomical boundaries do not necessarily correspond to underlying brain function. Therefore, regions of interest based on anatomical atlases display less signal homogeneity and so increase non-specific signal (Craddock et al., 2012). The second advantage of functional atlases is that no predetermined anatomical atlas is required during the analysis. Consequently, the independence of our pipeline provides the capacity to build brain networks in species that have not been fully investigated.

### 3.2. Modular organisation of mouse lemur brains

High modularity is an important principle of brain organisation (Bullmore and Sporns, 2009). It can be measured with modularity of a partition (Q). Here we found Q=0.43 in mouse lemurs. This value is consistent with Q values reported in rats (Q=0.39 (D’souza et al., 2014)), other non-human primates (0.33 < Q < 0.54 (Shen et al., 2012)) or humans in our study (Q = 0.56) and indicates that the mouse lemur brain can be partitioned into modules. Using graph analysis, we identified six cortical and subcortical modules that corresponded to large scale networks. This organisation into six modules is consistent with the number of modules reported in rats (n=6 (D’souza et al., 2014)), other non-human primates (for example n=4 (Shen et al., 2012) or n=7 (Grayson et al., 2016) in *Macaca fascicularis*), or humans in our study (n=6).

### 3.3. Characterisation of large scale networks in mouse lemur brains

#### 3.3.1. Multi-method approach of resting state analysis in animals

Whole brain networks can be decomposed into large scale networks. However, there are no absolute frontiers between these large scale networks due to the gradualness of the interactions between the different regions of the brain. Several methods, such as dictionary learning, graph analysis and seed-based studies can be used to identify these large scale networks in mammal brains. They rely on various mathematical bases associated with various sensitivities to image artefacts (Power et al., 2014). Also, these methods have diverse abilities to classify brain regions into networks. For example, graph analysis attributes each region to one and only one network while dictionary learning and seed-based analysis can attribute a region to several networks. In most resting state fMRI studies in animals, neuronal networks are identified on the basis of a single method. Here we showed that different methods do not detect exactly the same networks. However, networks identified with each method display a strong overlap. Functional regions included in a network by several methods represent the more robust parts of the network. Thus, we propose a first classification of the mouse lemur networks that takes into account only regions identified by two or three methods (Fig. 9, Suppl. Table 3). An overview of each network is presented in the following paragraphs.

**Figure 9.**
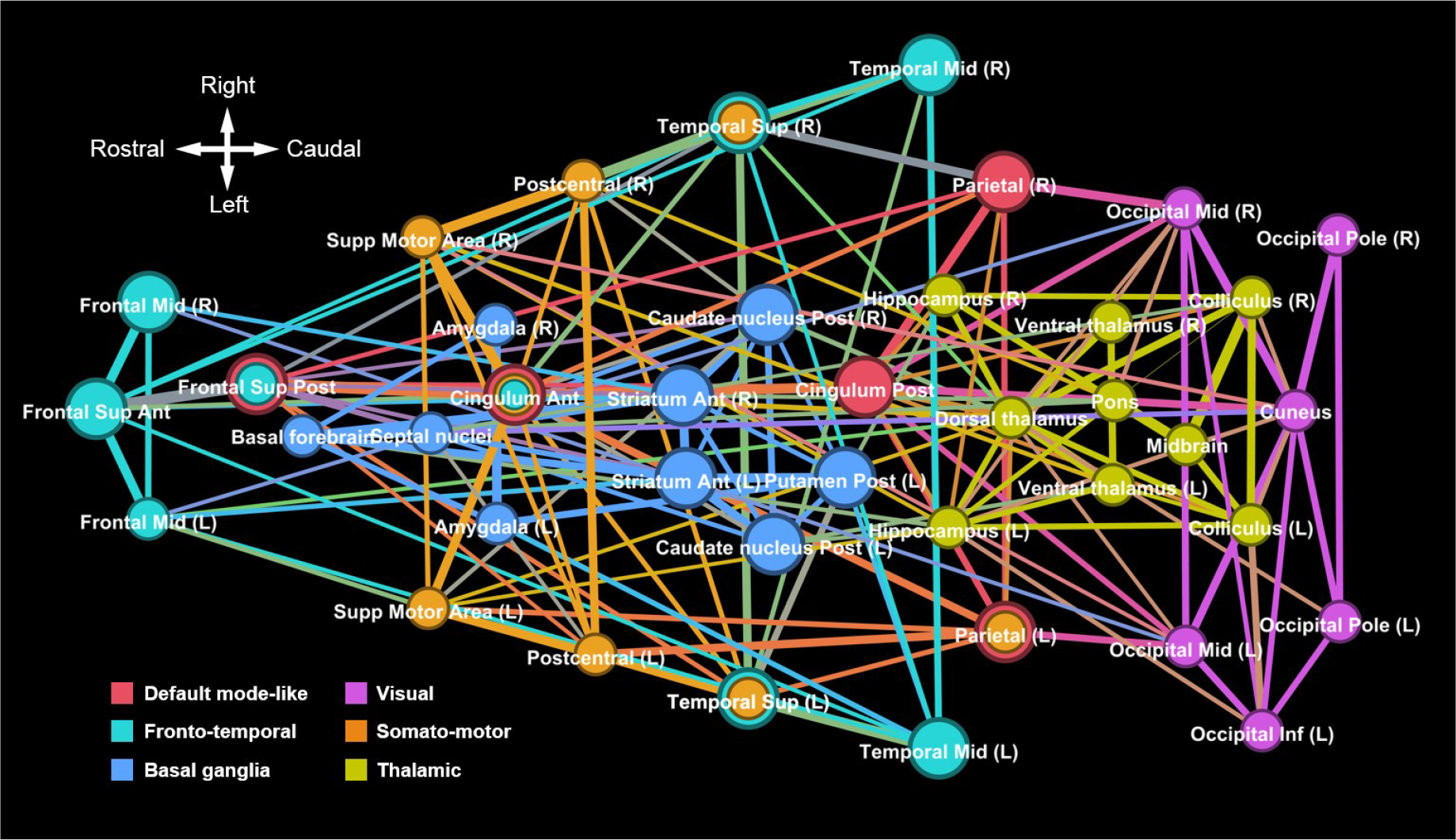
Most robust functional networks identified in mouse lemurs using two or three network identification methods. Regions from the DMN, visual, fronto-temporal, somato-motor, basal ganglia and thalamic networks that could be identified by two or three network identification methods are considered as robustly associated to a network and are displayed on this figure. For each network, edges were reported from those identified with graph analysis.

#### 3.3.2. Default-mode-like network

The DMN is one of the most studied networks in humans (Hampson et al., 2006) and other mammals including rodents (Lu et al., 2012) and non-human primates (J. L. Vincent et al., 2007). It plays a critical role in several physiological and pathological processes such as Alzheimer’s or Parkinson’s diseases (Buckner et al., 2005; Gao and Wu, 2016). In mouse lemurs, four regions of this network were detected with network identification methods: anterior and posterior cingulum cortices, superior posterior frontal cortex and parietal cortex. In several species, these regions are reported to be part of the default mode network (Belcher et al., 2013; J. L. Vincent et al., 2007).

In humans and other mammals, the DMN contains highly connected hub nodes. In the mouse lemur brain, we also found that it contained the most connected nodes. Given the importance of this network it was critical to characterize it in the mouse lemur, which is widely used as a model of neurodegenerative diseases (Kraska et al., 2011; Mestre-Frances et al., 2018).

#### 3.3.3. Fronto-temporal network

The fronto-temporal network was found in mouse lemurs with dictionary learning and seed-based analysis, but was split into two networks (frontal and temporal) with graph analysis. One of its components, the superior temporal cortex, was a strong hub in the mouse lemur brain. In primates, these regions are reported to be part of the executive network (Hutchison et al., 2012)

#### 3.3.4. Networks specialized in sensory and motor information processing

We also identified networks that could be classified as externally-driven. The first one is the visual network. It involved mainly occipital areas. This network has been described in numerous primates under task and rest conditions (Belcher et al., 2013). The second externally-driven network is the somato-motor network. It has also been widely defined in humans (Beckmann et al., 2005), primates (Nelissen and Vanduffel, 2011), and many other mammals (Sierakowiak et al., 2015). It integrates sensory input and motor commands. In mouse lemurs, we found that this network contains several hubs such as the anterior cingulum cortex, the superior motor area and the postcentral cortices.

#### 3.3.5. Subcortical networks

Finally, two networks were identified in subcortical areas. The first one involved the basal ganglia. Similar networks are described in primates (Belcher et al., 2013), and other mammals (Sierakowiak et al., 2015) and are involved in emotional, motivational, associative and cognitive functions (Herrero et al., 2002).

The second subcortical network involved several regions such as the ventral thalamus (a strong hub in mouse lemurs), dorsal thalamus, hippocampus, colliculus, pons and midbrain. It was called "thalamic network".

### 3.4. Small-worldness features of mouse lemur brains

We finally evaluated the small-worldness properties of the mouse lemur functional networks by calculating small-world coefficients σ and ω. Our results attested that mouse lemur networks have small-world properties (ω = 0.39). Interestingly, ω was much smaller in the human brain (ω = 0.08) than in the lemur brain suggesting stronger small-world properties in humans. The small-world configuration is considered as optimal for local information processing and for its global transfer. Indeed, small-world networks have the unique ability to have specialized regions while simultaneously exhibiting shared or distributed processing across all of the communicating regions of a network (Telesford et al., 2011).

### 3.5. Cross species comparison: homologies and divergence between humans and mouse lemur networks

In a last part of the study, cerebral networks were analyzed in humans with the same graph analysis and dictionary learning algorithms as the ones used in mouse lemurs. Two major differences were reported between the two species. First, large scale networks were only cortical in humans while they involved two subcortical networks in lemurs. Second, in humans, large scale networks involved more functional regions than in lemurs. This latter result is consistent with the stronger small-world organization in humans than in lemurs suggesting a better efficacy of whole brain networks in humans. These differences between the two species may be related to a better efficacy of neuronal networks in humans, but they could also be associated to different awareness levels as lemurs were anesthetized while humans were awake during image acquisition. Indeed, Barttfeld et al. compared connectivity measures in awake and anesthetized conditions in primates. They showed that under anesthesia, the more frequent functional connectivity patterns inherit the structure of anatomical connectivity and exhibit fewer small-world properties (Barttfeld et al., 2015).

Graph analysis revealed four similar modules (default mode-like, visual, frontal, and temporal networks) in mouse lemurs and humans, although their regional organization was not strictly identical. Two other modules detected in humans (somato-motor and temporo-insular) corresponded to networks that were not detected in lemurs. On the contrary, the two subcortical modules detected in lemurs (basal ganglia and thalamic networks) were not detected in humans. Because of the multiple regions involved in module description by graph analysis and because of the possibility to attribute a region to only one network with this method, it was difficult to further compare human and lemur networks with this technique.

Dictionary learning also revealed four similar networks (DMN, visual, fronto-temporal/supramarginal and somato-motor networks) in lemurs and humans (Table 2; Suppl. Fig. 4). In both species, the DMN network involved the cingulum, frontal, and parietal cortices. In mouse lemurs, it involved the superior posterior frontal cortex that was probably subdivided in two functional regions (frontal superior medial and frontal superior posterior cortices) in humans. Other regions such as the temporal cortex were included in the human DMN but not in the mouse lemur DMN. Interestingly, in both species, this network was the one in which highest hubness coefficients (eigenvectors) were detected. This reinforces the importance of this network for brain functional organization. In humans, the default mode network has been largely linked to self-referential thought, internal-oriented cognition and monitoring of the environment (Buckner et al., 2008). The strength and stability of this network in mouse lemurs under anaesthesia is consistent with the discovery of this network in many other anesthetized animals (J. L. Vincent et al., 2007). This suggests that it is an essential element of brain functional organization and that it may be dedicated to other tasks too.

In the visual network, occipital cortex was detected in both species. Additional more anterior-parietal regions such as the paracentral lobule and the postcentral were highlighted in humans. We cannot rule out that this wider extension in human dataset is not related to the wakefulness state as it induces a richer repertoire of functional configurations (Barttfeld et al., 2015).

In mouse lemurs, a network involving the anterior cingulum, frontal and temporal regions was classified as the fronto-temporal network. In humans, one network involving mostly the anterior cingulum and frontal regions could be homologous to this network. Interestingly, in lemurs, this networks also involved temporal (superior and medial temporal regions) while it involved parietal regions (supramarginal anterior and parietal inferior cortices) as well as additional regions (supplementary motor, cingulum median and opercular regions) in humans. This network could correspond to the control-executive network (Solé-Padullés et al., 2016). If the fronto-temporal network of mouse lemur is equivalent to the fronto-supramarginal human network, then this would suggest a shift of the functional region localized in the superior temporal area in lemurs towards a supramarginal location in humans.

The last comparable network was the somato-motor network. In humans it involved regions surrounding the central sulcus (precentral and postcentral regions) as well as the supplementary motor region. In lemurs, there is no central sulcus, but this network involved similar regions (precentral and postcentral regions) as well as the supplementary motor region. Interestingly, this part of the network seemed to have a more anterior position in the brain of lemurs than in humans. This is consistent with the more anterior part of the motor regions reported in lemurs by Le Gros Clark (Le Gros Clark, 1931) and Brodmann (Brodmann, 1999 (original in 1909)). This pattern is linked to the smaller size of the frontal region in lemurs as compared to humans. Finally, in humans, this region involved the paracentral and the precuneus anterior cortices while it involved the region classified as anterior cingulate cortex in the mouse lemur. These two regions are localized in the same area and we cannot rule out that the functional region classified as anterior cingulate cortex in lemur indeed involved the pre and post central lobule in addition to the anterior cingulate cortex.

### 3.6. Anaesthesia-related limitations

One of the objectives of this study was to describe for the first time neuronal networks in mouse lemurs. It was conducted on sedated animals using isoflurane with the lowest non-awakening isoflurane level possible for mouse lemurs (1.25%). Isoflurane is expected to decrease the functional connectivity but at high doses (superior to 1.5%) or after a long exposure (Hutchison et al., 2014; Li and Zhang, 2018). Evaluating resting state networks in anesthetised and not in awake animals is an obvious limitation of the study (Schroeter et al., 2014). However, several animal studies showed that the major functional networks are preserved under anaesthesia (J. L. Vincent et al., 2007). Here, we confirm this assumption by describing several networks, including a DMN-like in anesthetised mouse lemurs. In the future, one may also focus on resting state fMRI in awake mouse lemurs to possibly evaluate more physiological brain states and increase the number of nodes associated with each identified network. Such an approach is challenging but has already been performed in marmosets (Belcher et al., 2013) and macaques (Goense et al., 2008).

## 4. Conclusion

This study provides the first characterisation of functional brain networks in mouse lemur primates. Local functional regions were identified without using any anatomical atlas. Six large scale networks were identified using several complementary data-driven and hypothesis-based methods. Networks identified with each method displayed a strong overlap and we propose a first classification of the most robust mouse lemur networks by selecting only regions identified by two or three methods. We also proposed a second validation method by comparing networks in lemurs and human brains. Indeed, a strong homology was reported between well characterized human cortical networks and lemur cortical networks. This further suggests the accuracy of the identified mouse lemur networks. The mouse lemur brain displayed small-world features leading to optimal information transfer. Finally, critical hubs were detected and involved the posterior and anterior cingulate cortices, the central prefrontal cortex, and the dorsal thalamus.

The mouse lemur is an interesting primate because of its key position in the phylogenetic tree, rodent-like small size and nocturnal and arboreal lifestyle. The 3D functional atlas and resting state network maps are freely available at https://www.nitrc.org/projects/fmri_mouselemur/. The imaging tools used to create and manipulate the template are also available (https://sammba-mri.github.io).

## 5. Materials and methods

### 5.1. Animals and breeding

This study was carried out in accordance with the recommendations of the European Communities Council directive (2010/63/EU). The protocol was approved by the local ethics committees CEtEA-CEA DSV IdF (authorization 201506051736524 VI (APAFIS#778)). All mouse lemurs studied were born in the laboratory breeding colony of CNRS/MNHN in Brunoy, France (UMR 7179 CNRS/MNHN) and bred in our laboratory (Molecular Imaging Research Center, CEA, Fontenay-aux-Roses).

Sixteen mouse lemurs (12 males and 4 females) were initially included in this study. Two females that presented brain lesions on anatomical MRI were excluded from the analysis. The 14 analysed animals ranged from 0.9 to 3.1 years old (mean±SD: 1.7±0.7) (Suppl. Table 1). Housing conditions were cages containing one or two lemurs with jumping and hiding enrichment, temperature 24–26°C, relative humidity 55% and seasonal lighting (summer: 14 hours of light/10 hours of dark; winter: 10 hours of light/14 hours of dark). Food consisted of fresh apples and a homemade mixture of bananas, cereals, eggs and milk. Animals had free access to tap water. None of the animals had previously been involved in pharmacological trials or invasive studies.

### 5.2. Animal preparation and MRI acquisition

Each animal was scanned twice with an interval of 6 months. All scanning was under isoflurane anaesthesia at 1.25-1.5% in air, with respiratory rate monitored to confirm animal stability until the end of the experiment. Body temperature was maintained by an air heating system at 32°C, inducing a natural torpor in mouse lemurs (Aujard and Vasseur, 2001). This has the advantage of allowing a low anaesthesia level without reawakening.

The MRI system was an 11.7 Tesla Bruker BioSpec (Bruker, Ettlinger, Germany) running ParaVision 6.0.1. Anatomical images were acquired using a T2-weighted multi-slice multi-echo (MSME) sequence: TR = 5000 ms, TE = 17.5 ms, 6 echoes, inter-echo time = 5 ms, FOV = 32 × 32 mm, 75 slices of 0.2 mm thickness, resolution = 200 µm isotropic, acquisition duration 10 min. Resting state time series data were acquired using a gradient-echo EPI sequence: TR = 1000 ms, TE = 10.0 ms, flip angle = 90°, repetitions = 450, FOV = 30 × 20 mm, 23 slices of 0.9 mm thickness and 0.1 mm gap, resolution = 312 × 208 × 1000 µm, acquisition duration 7m30s.

### 5.3. MRI acquisition in humans

Forty-two healthy participants from the ‘Imagerie Multimodale de la Maladie d’Alzheimer à un stade Précoce’ (IMAP) study (Caen) were included in the present study (18 males and 24 females ranging from 41 to 60 years old (mean±SD: 50±5.9)). All participants were scanned on a 3.0 T scanner (Philips Achieva, Amsterdam, Netherlands) at the Cyceron Center (Caen, France). Anatomical T1-weighted images were acquired using a 3D fast-field echo sequence (3D-T1-FFE sagittal TR = 20 ms, TE = 4.6 ms, flip angle = 10°, 180 slices of 1 mm with no gap, FOV = 256 × 256 mm^2^, in-plane resolution = 1 × 1 mm^2^). Resting state time series data were acquired using an interleaved 2D T2* SENSE EPI (2D-T2*-FFE-EPI axial, SENSE = 2; TR = 2382 ms; TE = 30 ms; flip angle = 80°; 42 slices of 2.8 mm with no gap, repetitions = 450, FOV = 224 × 224 mm^2^, in plane resolution = 2.8 × 2.8 mm^2^, acquisition duration = 11.5 min). Head motion was minimized with foam pads. Participants were equipped with earplugs and the scanner room’s light was turned off. During this acquisition, participants were asked to keep their eyes closed and relax without falling asleep.

### 5.4. MRI pre-processing

#### 5.4.1. Mouse lemur data

Scanner data were exported as DICOM files then converted into NIfTI-1 format. Then spatial pre-processing was performed using the python module sammba-mri (SmAll MaMmals BrAin MRI; http://sammba-mri.github.io) which, using nipype for pipelining (Gorgolewski et al., 2011), leverages AFNI (Cox, 1996) for most steps and RATS (Oguz et al., 2014) for brain extraction. Anatomical images were mutually registered to create a study template, which was further registered to a high resolution anatomical mouse lemur template (Nadkarni et al., 2018). Resting state images were corrected for slice timing (interleaved), motion, and B0 distortion (per-slice registration to respective anatomicals), then all brought into the same space of the mouse lemur template by successive application of the individual anatomical to study template and study template to mouse lemur atlas transforms. Functional images were further pretreated using Nilearn (Abraham et al., 2014). Nuisance signal regression was applied including a linear trend as well as 24-motion confounds (6 motion parameters, those of the preceding volume, plus each of their squares (Friston et al., 1994)). Images were then spatially smoothed with a 0.9 mm full-width at half-maximum Gaussian filter. The first 10 volumes were excluded from analysis to ensure steady-state magnetization.

#### 5.4.2. Human data

Artefacts were inspected in individual datasets using the TSDiffAna routines (http://imaging.mrc-cbu.cam.ac.uk/imaging/DataDiagnostics). Datasets displaying significant movements (> 1.5° rotation or > 3 mm translation) and abnormal variance distribution and/or artefacted were excluded from the analysis. Data were then preprocessed as defined in Landeau *et al.* (Landeau et al., 2017) with slice timing correction, realignment to the first volume and spatial normalization within native space to correct for distortion effects. EPI volumes were registered to their own high resolution anatomical image and then registered and normalized to MNI template space. Nuisance signal regression was applied including a linear trend as well as 24-motion confounds (6 motion parameters, those of the preceding volume, plus each of their squares (Friston et al., 1994)). Images were then spatially smoothed with a 2 mm full-width at half-maximum Gaussian filter.

### 5.5. Identification of functional regions by dictionary learning and creation of a 3D functional atlas

Multi-animal dictionary learning was performed with Nilearn (Mensch et al., 2016) on preprocessed resting state functional MR images. A mask excluding the corpus callosum, hindbrain, ventricles and three systematically artefacted regions (olfactory bulb, ventral entorhinal cortex and prepiriform cortex) was used to restrict functional data to non-noise voxels prior to dictionary learning analysis. During a pilot investigation, several analyses were performed using 20, 30, 35, 40, 45, 50, and 60 sparse components (SCs). The study based on 35 SCs was selected for the final analysis as it highlighted either unilateral local functional regions or bilateral regions. Moreover, the extracted components matched well to anatomy (Nadkarni et al., 2018). The 35 SCs were used to create a 3D functional atlas of the mouse lemur brain. Each bilateral SC was split into two unilateral regions. Regions smaller than 5 mm^3^ were excluded leading to 48 local functional regions. Each region was then named using ITK-SNAP to create a 3D functional atlas (Yushkevich et al., 2006). The same procedure than in lemurs was applied to process human fMRI data. We used 35 SCs and a grey matter mask without hindbrain.

### 5.6. Identification of large scale networks

#### 5.6.1. Connectivity matrix based on functional atlas

Partial correlation matrices were created using fully preprocessed MR images by calculating the partial correlation coefficients between BOLD MR signal timecourses within each region of the 3D functional atlas. Partial correlations were used because they select direct associations between regions and allow the control of indirect correlations (Mechling et al., 2014). Individual partial correlation matrices were computed from shrunk covariance matrices using the Ledoit and Wolf shrinkage coefficient (Ledoit and Wolf, 2004) as recommended by Varoquaux et al. (Varoquaux et al., 2012) and Brier et al. (Brier et al., 2015). Partial correlation coefficients were then Fisher’s *z*-transformed. Values from different animals were averaged and thresholded based on a one-tailed *t*-test (*p* ≤ 0.01) (Mechling et al., 2014).

#### 5.6.2. Modularity and large scale network identification by graph theory analysis

The modularity of a partition (Q) is the degree to which a network can be subdivided into non-overlapping groups of nodes (D. B. Vincent et al., 2008). The modularity of a partition as well as an optimal segregation of the whole brain network into modules were calculated using Gephi 0.9.2 (Bastian et al., 2009).

#### 5.6.3. Large scale network identification by dictionary learning analysis

A second dictionary analysis was performed in mouse lemurs and humans using a smaller number of SCs in order to highlight large networks and to compare them. Six SCs were used based on the 6 modules found with the graph theory analysis (see Results). In humans, a mask excluding the hindbrain and the white matter was used prior to the analysis to compare the dictionary learning of the two species in a similar space.

#### 5.6.4. Large scale network identification by seed-based analysis

Seeds corresponded to each region of the 3D functional atlas. The BOLD signal was averaged within each seed. The functional connection between the seed’s mean BOLD signal and the BOLD signal in any voxel of the brain was estimated using a first-level general linear model (Nistats (Abraham et al., 2014)). The within-animal effect (*i.e.* the two series of MR images from each animal) was entered as a predictor (design matrix) and the mean seed time course as regressor. The model directly returned a fixed effect of the seed across the two sessions, producing 14 *z*-statistic maps. The functional regions previously identified were used as seeds. For each seed, a visual inspection of the animal mean *z*-statistic maps allowed the selection of four distinct large scale networks that were spread over the whole brain.

### 5.7. Identification of functional regions from dictionary-learning and seed-based maps

Dictionary learning and seed-based analysis produced maps showing pixels belonging to different networks. These maps were extracted and pasted into the 3D functional atlas. A brain region was considered to be part of a specific network when the volume of labelled voxels within the map occupied at least 30% of that region.

### 5.8. Evaluation of functional hubness and small-worldness features of mouse lemur brains by graph theory analysis

We consider in this analysis the absolute value of the correlation coefficient as performed routinely in human fMRI graph theory studies (De Vico Fallani et al., 2014).

#### 5.8.1. Brain hubs in mouse lemurs

Eigenvector centrality, a measure of "hubness", was measured using NetworkX (Hagberg et al., 2008).

#### 5.8.2. Small-worldness of mouse lemur brain networks

Network topology can be characterized using two small-world coefficients (σ and ω) (NetworkX (Hagberg et al., 2008)).

σ is defined as 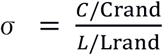 (Watts and Strogatz, 1998)
ω is defined as 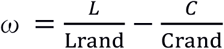(Telesford et al., 2011).

With C and L being, respectively, the average clustering coefficient (a measure of network segregation) and the average shortest path length (a measure of integration) of the network. Crand and Lrand are their equivalent derived random networks. Small-world networks have σ values superior to 1 and ω values close to 0 (Telesford et al., 2011).

## 6. Acknowledgements

We thank the France-Alzheimer Association, Plan Alzheimer Foundation, Neuratris and the French Public Investment Bank’s “ROMANE” program for funding this study. We thank all the persons who contributed to IMAP+ data acquisition and analyses: Florence Mézenge, Renaud La Joie, Julie Gonneaud, Audrey Perrotin, Robin de Flores, Clémence Tomadesso, Justine Mutlu, Nicolas Villain, Marine Fouquet, Katell Mevel, Francis Eustache, Béatrice Desgranges, Stéphanie Egret, Vincent de La Sayette, Fausto Viader, Alice Pélerin, Malo Gaubert, Géraldine Poisnel, Anne Quillard, Ahmed Abbas, Louisa Barré, Alain Manrique, Florence Pasquier, the Cyceron staff members; and the volunteers who were included in this study.

