## Supplementary material for "Resting state cerebral networks in mouse lemur primates: from multilevel validation to comparison with humans": Suppl.

Supplementary Data

### Supplementary Tables

| | *Sex* | *Age (months)* | *Age (years)* | *Anatomical brain lesion* |
|---|---|---|---|---|
| **283EA** | M | 10.6 | 0.9 | No |
| **365A** | M | 10.6 | 0.9 | No |
| **285AB** | M | 10.7 | 0.9 | No |
| **285AAA** | M | 16.5 | 1.4 | No |
| **283CCA** | M | 16.6 | 1.4 | No |
| **263BCE** | M | 17.8 | 1.5 | No |
| **314CA** | M | 18.0 | 1.5 | No |
| **283CA** | M | 22.4 | 1.9 | No |
| **285E** | M | 22.6 | 1.9 | No |
| **276BC** | M | 28.0 | 2.3 | No |
| **285D** | M | 28.1 | 2.3 | No |
| **289BB** | F | 28.8 | 2.4 | No |
| **300BA** | M | 29.8 | 2.5 | No |
| ***288BC*** | F | 37.3 | 3.1 | Yes |
| **208CBF** | F | 37.5 | 3.1 | No |
| ***310C*** | F | 39.9 | 3.3 | Yes |

**Supplementary Table 1. Cohort of mouse lemurs involved in the study.**

<table>
<tr><th></th><th>Modules</th><th>Dictionary learning</th><th>Seed-based</th></tr>
<tr><td rowspan="6">Default mode-like</td><td colspan="3">Cingulum Post</td></tr>
<tr><td colspan="3">Cingulum Ant</td></tr>
<tr><td colspan="3">Frontal Sup Post</td></tr>
<tr><td colspan="3">Parietal (R&L)</td></tr>
<tr><td>Supp Motor Area (R&L)</td><td rowspan="2"></td><td>Frontal Mid</td></tr>
<tr><td>Postcentral (R&L)</td><td>Dorsal thalamus</td></tr>
<tr><td rowspan="5">Visual</td><td colspan="2">Occipital Pole (R&L)</td><td rowspan="5"></td></tr>
<tr><td colspan="2">Cuneus</td></tr>
<tr><td colspan="2">Occipital Mid (R&L)</td></tr>
<tr><td colspan="2">Occipital Inf (L)</td></tr>
<tr><td>Temporal Inf (L)</td><td>Occipital Inf (R)</td></tr>
<tr><td rowspan="5">Frontal</td><td colspan="3">Frontal Sup Ant</td></tr>
<tr><td colspan="2">Frontal Mid (R&L)</td><td></td></tr>
<tr><td>Precentral (R&L)</td><td colspan="2">Frontal Sup Post</td></tr>
<tr><td></td><td colspan="2">Cingulum Ant</td></tr>
<tr><td colspan="3"></td></tr>
<tr><td rowspan="4">Temporal</td><td colspan="3">Temporal Sup (R&L)</td></tr>
<tr><td colspan="3">Temporal Mid (R&L)</td></tr>
<tr><td>Temporal Inf (R)</td><td colspan="2" rowspan="2"></td></tr>
<tr><td>Posterior putamen (R)</td></tr>
<tr><td rowspan="11">Somato-motor</td><td rowspan="11"></td><td colspan="2">Supp Motor Area (R&L)</td></tr>
<tr><td colspan="2">Postcentral (R&L)</td></tr>
<tr><td colspan="2">Temporal Sup (R&L)</td></tr>
<tr><td colspan="2">Cingulum Ant</td></tr>
<tr><td colspan="2">Parietal (L)</td></tr>
<tr><td>Precentral (R&L)</td><td>Parietal (R)</td></tr>
<tr><td rowspan="5"></td><td>Frontal Mid</td></tr>
<tr><td>Frontal Sup Post</td></tr>
<tr><td>Cingulum Post</td></tr>
<tr><td>Caudate nucleus Post (R)</td></tr>
<tr><td>Dorsal thalamus</td></tr>
<tr><td rowspan="8">Basal ganglia</td><td colspan="3">Striatum Ant (R&L)</td></tr>
<tr><td colspan="3">Caudate nucleus Post (R&L)</td></tr>
<tr><td colspan="3">Putamen Post (L)</td></tr>
<tr><td colspan="2">Amygdala (R&L)</td><td>Frontal Sup Post</td></tr>
<tr><td colspan="2">Basal forebrain</td><td>Cingulum Ant</td></tr>
<tr><td colspan="2">Septal nuclei</td><td rowspan="3"></td></tr>
<tr><td>Hypothalamus</td><td>Putamen Post (R)</td></tr>
<tr><td>Globus pallidus</td><td></td></tr>
<tr><td rowspan="9">Thalamic</td><td colspan="2">Dorsal thalamus</td><td rowspan="9"></td></tr>
<tr><td colspan="2">Ventral thalamus (R&L)</td></tr>
<tr><td colspan="2">Hippocampus (R&L)</td></tr>
<tr><td colspan="2">Colliculus (R&L)</td></tr>
<tr><td colspan="2">Pons</td></tr>
<tr><td colspan="2">Midbrain</td></tr>
<tr><td>Occipital Inf (R)</td><td>Basal forebrain</td></tr>
<tr><td rowspan="2"></td><td>Septal nuclei</td></tr>
<tr><td>Globus pallidus</td></tr>
</table>

**Supplementary Table 2. Comparison of the regions belonging to the different networks extracted with module, dictionary learning and seed-based analysis.** Regions that were identified with different methods are grouped within a single case. The 3D functional atlas was pasted on different networks obtained by dictionary learning or seed-based analysis. A region was considered to belong to a network when more than 30% of its volume belonged to this network.

| | Robust functional regions |
|---|---|
| **Default mode-like** | Cingulum Post |
| | Cingulum Ant |
| | Frontal Sup Post |
| | Parietal (R&L) |
| **Visual** | Occipital Pole (R&L) |
| | Occipital Mid (R&L) |
| | Occipital Inf (L) |
| | Cuneus |
| **Fronto-temporal** | Frontal Sup Ant |
| | Frontal Mid (R&L) |
| | Frontal Sup Post |
| | Cingulum Ant |
| | Temporal Sup (R&L) |
| | Temporal Mid (R&L) |
| **Somato-motor** | Postcentral (R&L) |
| | Supp Motor Area (R&L) |
| | Temporal Sup (R&L) |
| | Cingulum Ant |
| | Parietal (L) |
| **Basal ganglia** | Striatum Ant (R&L) |
| | Caudate nucleus Post (R&L) |
| | Putamen Post (L) |
| | Amygdala (R&L) |
| | Basal forebrain |
| | Septal nuclei |
| **Thalamic** | Dorsal thalamus |
| | Ventral thalamus (R&L) |
| | Hippocampus (R&L) |
| | Colliculus (R&L) |
| | Pons |
| | Midbrain |

**Supplementary Table 3. Robust functional networks in mouse lemur primates.** This table displays components that could be identified by two or three methods. The different networks are the DMN, visual, fronto-temporal, somato-motor, basal ganglia and thalamic networks.

### Supplementary Figures

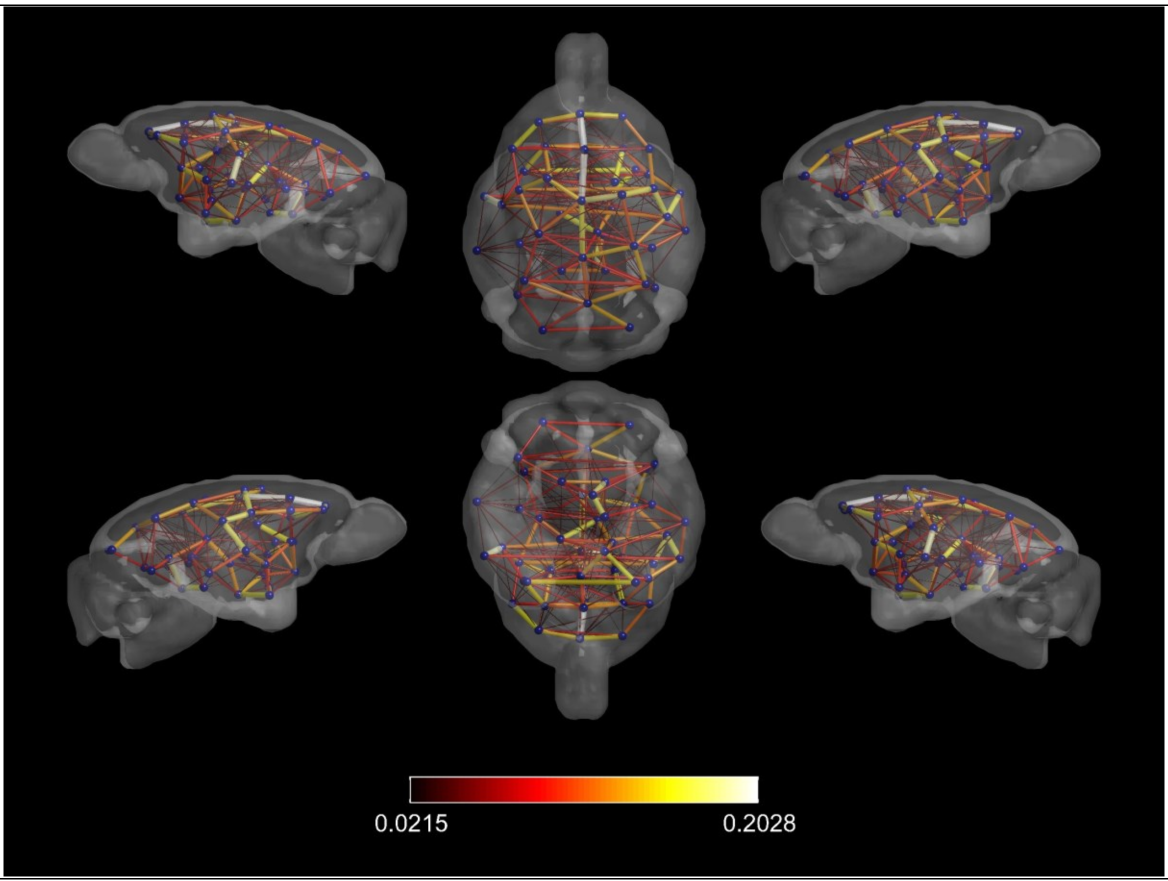


**Supplementary Figure 1. Whole brain network in mouse lemurs.**
Mean correlation matrix showing the mouse lemur brain network represented on a 3D mouse lemur brain space using BrainNet (Xia et al., 2013). Nodes represent the local functional regions extracted from our 3D functional atlas. They were spatially distributed based on their center of mass. Edges between the nodes represent the mean partial correlation from the 28 mouse lemurs. Color and size of these edges are proportional to this correlation. The color bar represents partial correlation values.

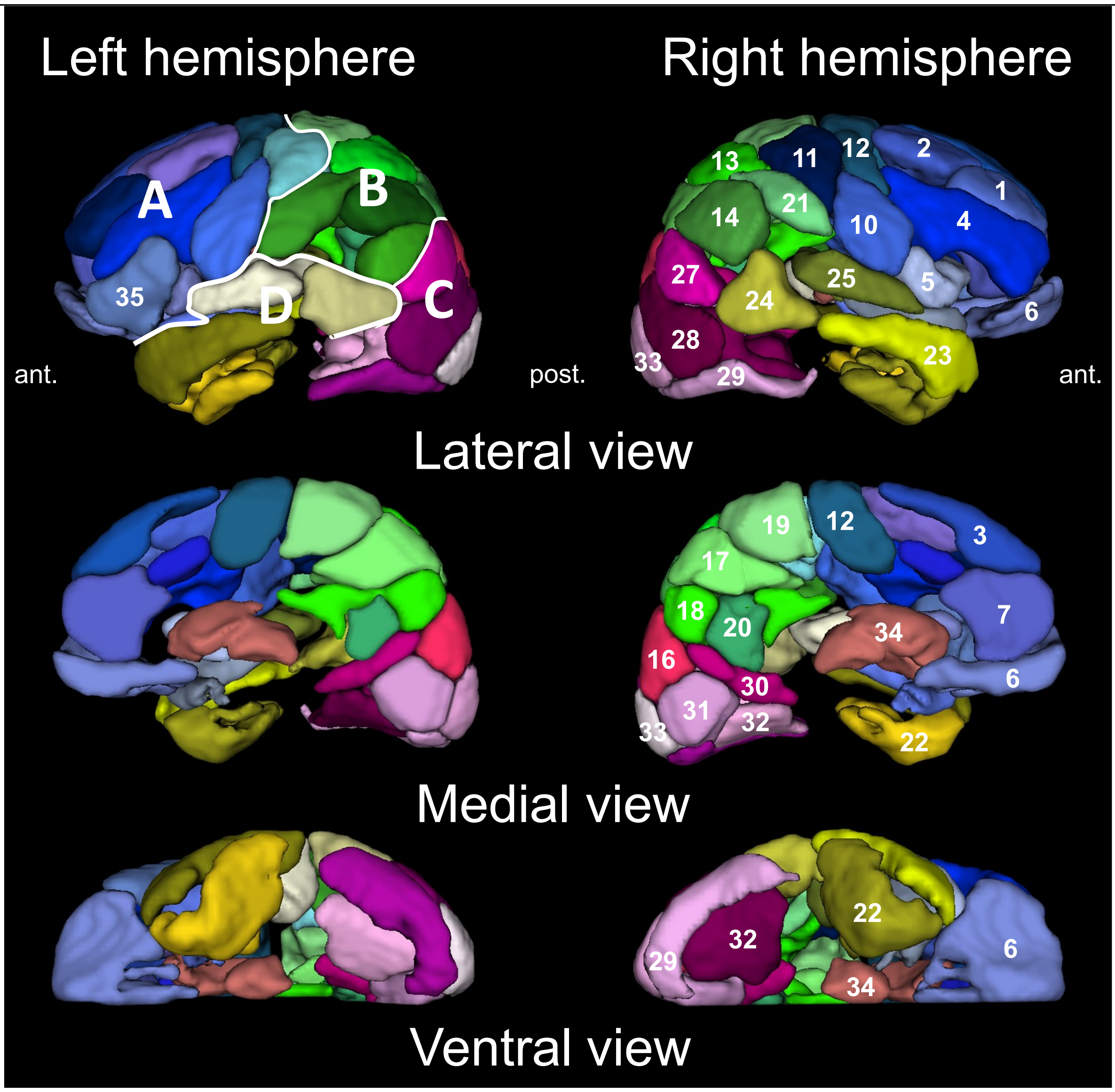


**Supplementary Figure 2. Human 3D functional atlas based on dictionary learning.**

Functional regions based on dictionary learning. Fifty six local functional regions were identified from the 35 sparse components (region volume ≥ 5000 mm$^3$). Brain regions were classified based on their locations within the frontal (A), parietal (B), occipital (C), and temporal (D) lobes and on the "AAL for SPM12" atlas (Tzourio-Mazoyer et al., 2002). We display three different views. 1. Frontal Superior Anterior, 2. Frontal Superior Posterior, 3. Frontal Superior Medial, 4. Frontal Middle, 5. Frontal Inferior Opercular, 6. Frontal Orbital, 7. Cingulum Anterior, 8. Cingulum Middle, 9. Insula, 10. Precentral, 11. Postcentral, 12. Supplementary Motor Area, 13. Parietal Superior Anterior, 14. Parietal Inferior, 15. Angular, 16. Parietal Superior Posterior, 17. Precuneus Anterior, 18. Precuneus Posterior, 19. Paracentral Lobule, 20. Cingulum Posterior, 21. Supramarginal, 22. Temporal Inferior, 23. Temporal Middle Anterior, 24. Temporal Middle Posterior, 25. Temporal Superior, 26. Cuneus, 27. Occipital Superior, 28. Occipital Middle, 29. Occipital Inferior, 30. Calcarine, 31. Lingual, 32. Fusiform, 33. Occipital Pole, 34. Striatum-Thalamus, 35. Frontal Inferior.

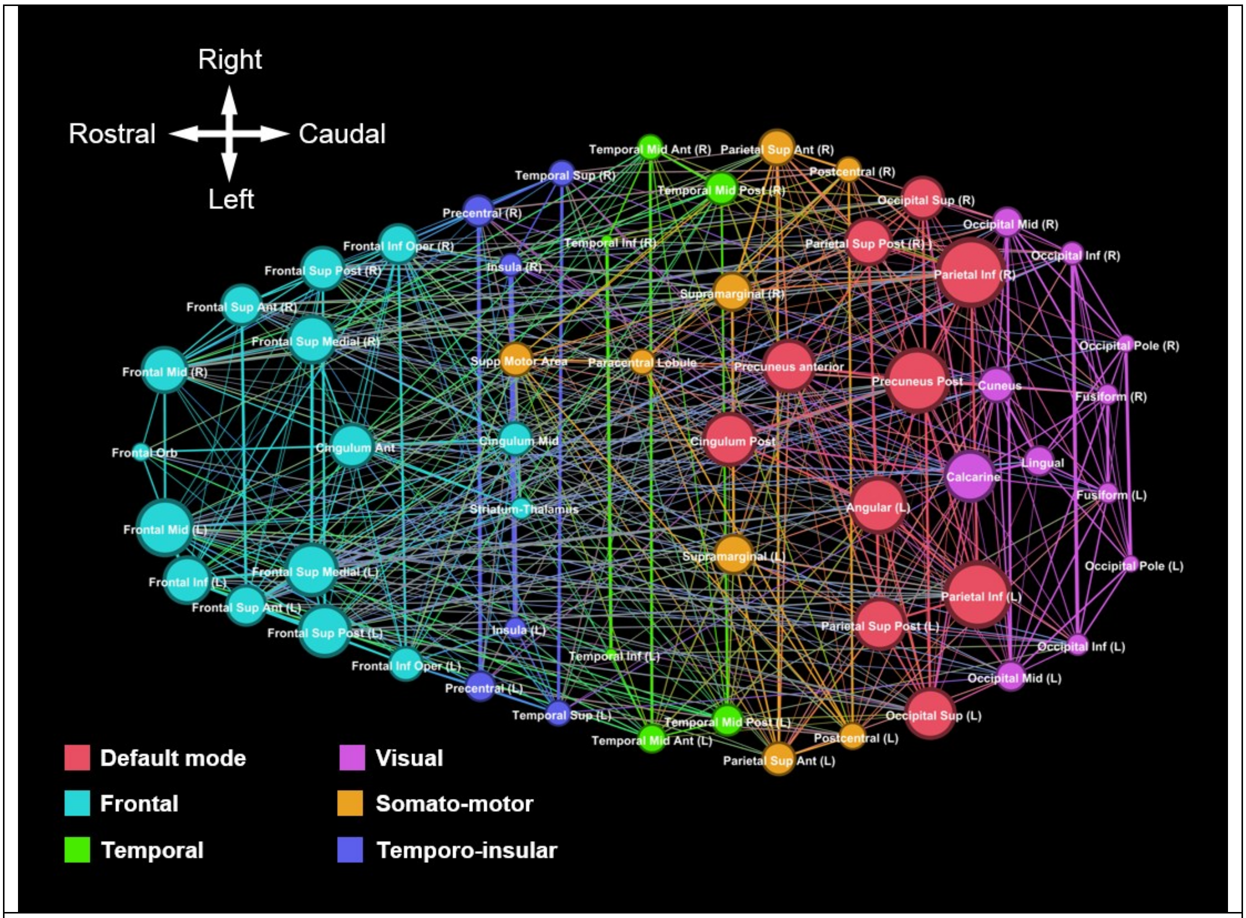


**Supplementary Figure 3. Human networks identified using graph analysis based on 56 functional regions.**

Using graph analysis, we partitioned the human brain into six cortical and subcortical modules. A color and a name were assigned to each module. Colors highlight interactions between different nodes, *i.e.* they outline large scale networks. Eigenvector centrality, a measure of node influence, is represented by the node size.

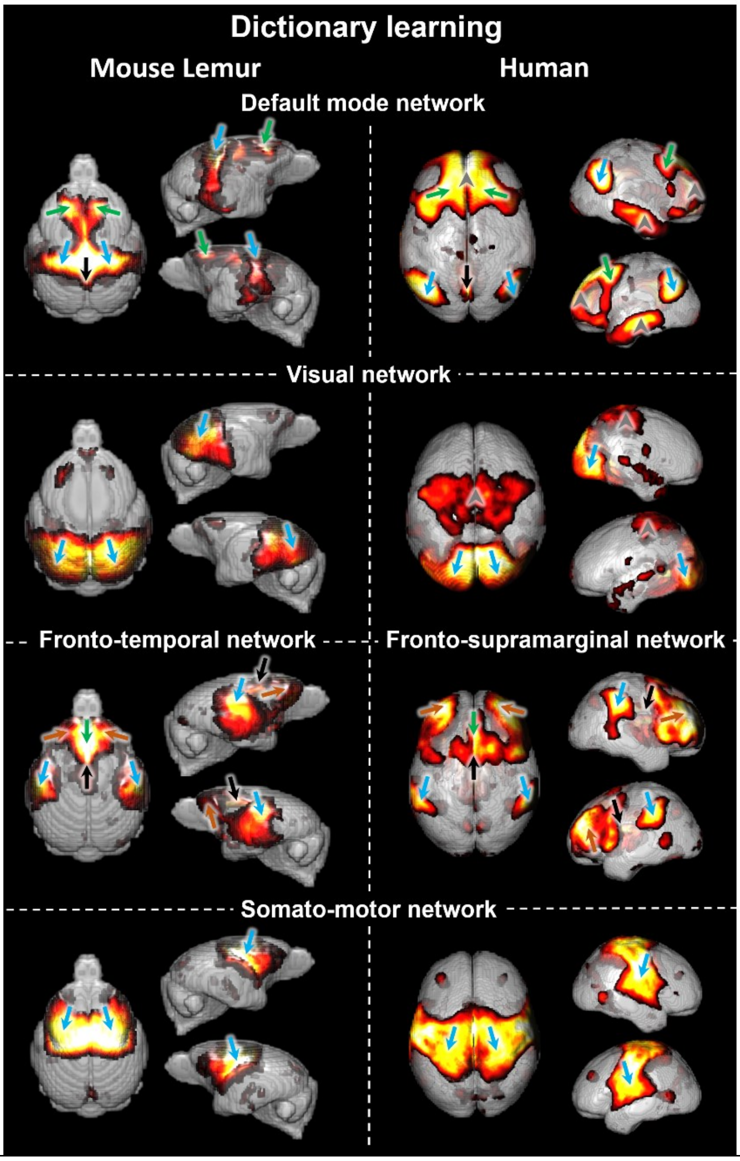


**Supplementary Figure 4. Comparison of the resting state network organization in humans and mouse lemurs.**

Functional spatial maps extracted with dictionary learning are displayed side by side. Four cortical networks were matched between lemurs and humans. They were classified as default mode network, visual, fronto-temporal/supramarginal, and somato-motor. Regions that are similar across species are pointed out with arrows. In the DMN-like network, frontal cortex (green arrows), posterior cingulum cortex (black arrows), parietal cortex (blue arrows) were detected in both species. The superior medial frontal and temporal cortices (arrow-heads) were detected in humans but not in mouse lemurs. In the visual network, occipital cortex was detected in both species. An additional independent region was detected in the paracentral lobule and postcentral cortices in humans (arrow-head). For the fronto-temporal/supramarginal network the middle frontal (brown arrows), superior frontal (green arrows) and anterior cingulate cortex (black arrow) were detected in both species. Interestingly, in humans the supramarginal cortex seems to fit with the superior temporal cortex, in lemurs (blue arrows). For the somato-motor network, regions were detected on both side of the central sulcus in humans and in a similar region in lemurs (in which there is no central sulcus). In humans, they were in parieto-frontal regions while in lemurs they involved more frontal regions (blue arrows).
